# Sex stratified analyses enable new genetic insights into brain imaging phenotypes

**DOI:** 10.64898/2026.04.21.719541

**Authors:** Nannan Zhang, Shaoying Wang, Jilian Fu, Yuan Ji, Nana Liu, Qian Qian, Hui Xue, Hao Ding, Meng Liang, Wen Qin, Jiayuan Xu, Chunshui Yu

**Author notes:** Correspondence to: Chunshui Yu, Jiayuan Xu, Wen Qin. These authors contributed equally to this work: Nannan Zhang, Shaoying Wang, Jilian Fu, Yuan Ji.

## Abstract

Sex differences are commonly observed in neuroimaging phenotypes and in the risk of brain diseases, yet the underlying genetic mechanisms remain poorly understood. We investigated sex differences in the genetic architecture of 805 neuroimaging phenotypes in 22,950 males and 22,950 females matched for sample size and covariates, and systematically compared sex-stratified with sex-combined genetic analyses. We found eight variant-trait associations with significant sex differences, 235 fine-mapped sex-dominant causal associations, 457 sex-dominant colocalizations with sex hormones, and 96 sex-dominant colocalizations with schizophrenia. Compared with sex-combined analysis, sex-stratified analysis identified 47 new genetic associations, 170 new fine-mapped causal associations, 1,019 new colocalizations with sex hormones, and 191 new colocalizations with schizophrenia. Additionally, sex-stratified analysis improved global heritability and genetic-correlation estimates and enhanced polygenic prediction for certain phenotypes. This work highlights the need to routinely perform sex-stratified genetic association analyses to elucidate sex-specific and sex-shared genetic control of neuroimaging phenotypes and related disorders.

## Introduction

Sex differences are commonly observed in human brain anatomy, as measured *in vivo* by imaging-derived phenotypes (IDPs), including gray matter volume (GMV), cortical thickness (CT), surface area (SA), and white matter microstructure^1–5^. They may arise from differing genetic architectures, environmental pressures, neurodevelopment, and hormone milieu between men and women^6–8^, and contribute to sex differences in brain functional activity and connectivity, cognitive performance, emotion regulation, and the epidemiology of neuropsychiatric disorders^9–12^. Although brain anatomical IDPs are highly heritable^13,14^, the genetic architecture underlying their sex differences remains poorly understood, because most previous genome-wide association studies (GWASs) of brain anatomical IDPs were conducted on sex-combined samples^15–17^, which may be biased by sex differences in genetic effects^6,18^. Identifying the genetic architecture underlying sex differences in brain anatomy is crucial for understanding the biological mechanisms driving sex disparities in brain-related traits, including disorders, and for developing personalized treatments.

Two previous studies examined sex differences in the genetic architecture of brain IDPs via sex-stratified GWASs. One reported sex differences in two single nucleotide polymorphism (SNP) effects on amygdala subregional volumes^18^. Another assessed sex differences in SNP-based heritability estimates, genetic correlations, and SNP effects for 1,106 GMV, CT, and SA phenotypes^6^. That study reported an overall similar genetic architecture of these IDPs between sexes but revealed significant sex differences in mean heritability for GMV-IDPs and SA-IDPs, genetic correlations for two IDPs, and SNP effects for two other IDPs^6^. Although these studies advanced understanding of sex differences in the genetic architecture of brain IDPs, several limitations remain. First, they compared males and females who were not matched for sample size and covariates, which may introduce bias. Second, they focused exclusively on brain gray matter IDPs, leaving sex differences in the genetic architecture of brain white matter IDPs uncertain. Third, other genetic characteristics remain unexplored. For instance, statistical fine-mapping can prioritize causal variant-trait associations for functional experiments, and colocalization analysis can identify shared genetic architecture between brain IDPs and other brain-related traits. However, sex differences in causal associations and genetic colocalizations remain unclear.

Compared with sex-combined GWAS, sex-stratified GWAS can not only identify sex differences in genetic architecture, but also reveal new genetic insights and improve the precision of genetic analyses and their clinical applications. For example, a previous study on non-imaging phenotypes indicates that sex-stratified analyses can discovery novel trait-associated loci and improve polygenic score (PGS) prediction of those traits^19^. However, a systematic comparison between sex-stratified and sex-combined GWASs regarding their performance in heritability estimation, genetic correlation, association discovery, PGS prediction, statistical fine-mapping, and genetic colocalization is still lacking. For instance, colocalization analysis using sex-stratified GWASs may reveal sex-dominant colocalizations with sex hormones and brain-related disorders that show sex-preferred epidemiology (e.g., higher male-to-female prevalence in schizophrenia).

Based on 48,781 participants (23,060 males and 25,721 females) recruited by the UK Biobank (UKB)^20^, we created male and female subgroups matched for sample size (n = 22,950) and covariates, along with a sex-combined group (n = 45,900) by including participants from both subgroups. We focused on 805 brain IDPs reflecting gray matter morphology or white matter integrity, and systematically investigated sex differences in heritability estimates, genetic correlations, SNP effects, causal associations, and colocalizations. In the same sample (n = 45,900), we also compared sex-stratified and sex-combined genetic analyses for their performance in heritability estimation, genetic correlation, association discovery, PGS prediction, statistical fine-mapping, and genetic colocalization. The overall workflow is illustrated in Extended Data Fig. 1.

## Results

### Participants

All participants were recruited from the UK Biobank^20^. After quality control of genetic and neuroimaging data, we included 48,781 participants (23,060 males and 25,721 females). The demographic and covariate data are shown in Supplementary Table 1. We conducted propensity score matching (PSM)^21^ to create male and female subgroups that were matched in sample size (n = 22,950) and covariates. After matching, the absolute standardized mean differences (SMDs) in all covariates between the two sex subgroups were less than 0.1 (Extended Data Fig. 2; Supplementary Table 2), indicating good balance. The 22,950 males and 22,950 females together comprised the sex-combined sample (n = 45,900). We performed genetic analyses on 16,013,685 autosomal variants and 805 brain IDPs derived from structural (sMRI) and diffusion (dMRI) magnetic resonance imaging data (Supplementary Table 3). The 373 sMRI-IDPs comprised the GMVs of 139 brain regions from volume-based segmentation, the CTs and SAs of 62 cortical regions from surface-based segmentation, and the volumes of 110 subcortical subregions from Freesurfer subsegmentation. The 432 dMRI-IDPs evaluated nine white matter integrity measures across 48 tracts.

### Heritability estimation

SNP-based heritability estimates the proportion of variance in a trait explained by the additive effects of genetic variants within a given sample. Sex difference in heritability suggests variation in the genetic mechanisms underlying the trait in males and females. We utilized LDSC v.1.0.0^22^ to estimate SNP-based heritability for all IDPs in males, females, and sex-combined participants, respectively (Supplementary Table 4).

#### Sex difference in heritability estimates

We performed three analyses to evaluate sex differences in heritability estimates of the 805 IDPs. First, we calculated Spearman’s rank correlation of heritability estimates of IDPs between males and females, revealing strong but not perfect correlations across all IDPs (*rho* = 0.90, *P* = 2.41 × 10^-285^; Fig. 1a), sMRI-IDPs (*rho* = 0.90, *P* = 2.63 × 10^-135^; Extended Data Fig. 3a), and dMRI-IDPs (*rho* = 0.88, *P* = 4.09 × 10^-140^; Extended Data Fig. 3b). Second, we conducted a paired-sample Wilcoxon signed-rank test to assess overall sex difference in heritability of these IDPs, revealing higher heritability in females than in males (*P* = 1.31 × 10^-10^; median *h^2^* difference = 0.0075; Fig. 1b). The greater female-to-male heritability was also observed for both sMRI-IDPs (*P* = 2.33 × 10^-4^; median *h^2^* difference = 0.0064) and dMRI-IDPs (*P* = 6.88 × 10^-8^; median *h^2^* difference = 0.009) (Extended Data Fig. 3c). Third, we performed a *Z*-test to identify sex difference in heritability estimates for each IDP. Although no differences remained significant at *P* < 0.05/805 = 6.21 × 10^-5^ (Bonferroni corrected), 24 brain IDPs showed sex differences in heritability at *P* < 0.05 (Fig. 1a and Supplementary Table 5), with the most pronounced sex difference (*h^2^_male_* = 0.25, *h^2^_female_* = 0.14, *Z* = 2.79, *P* = 5.34 × 10^-3^) in L1 in the anterior limb of the right internal capsule. The subtle sex differences may account for the overall high correlation yet differing heritability estimates of IDPs between males and females.

**Fig. 1.**
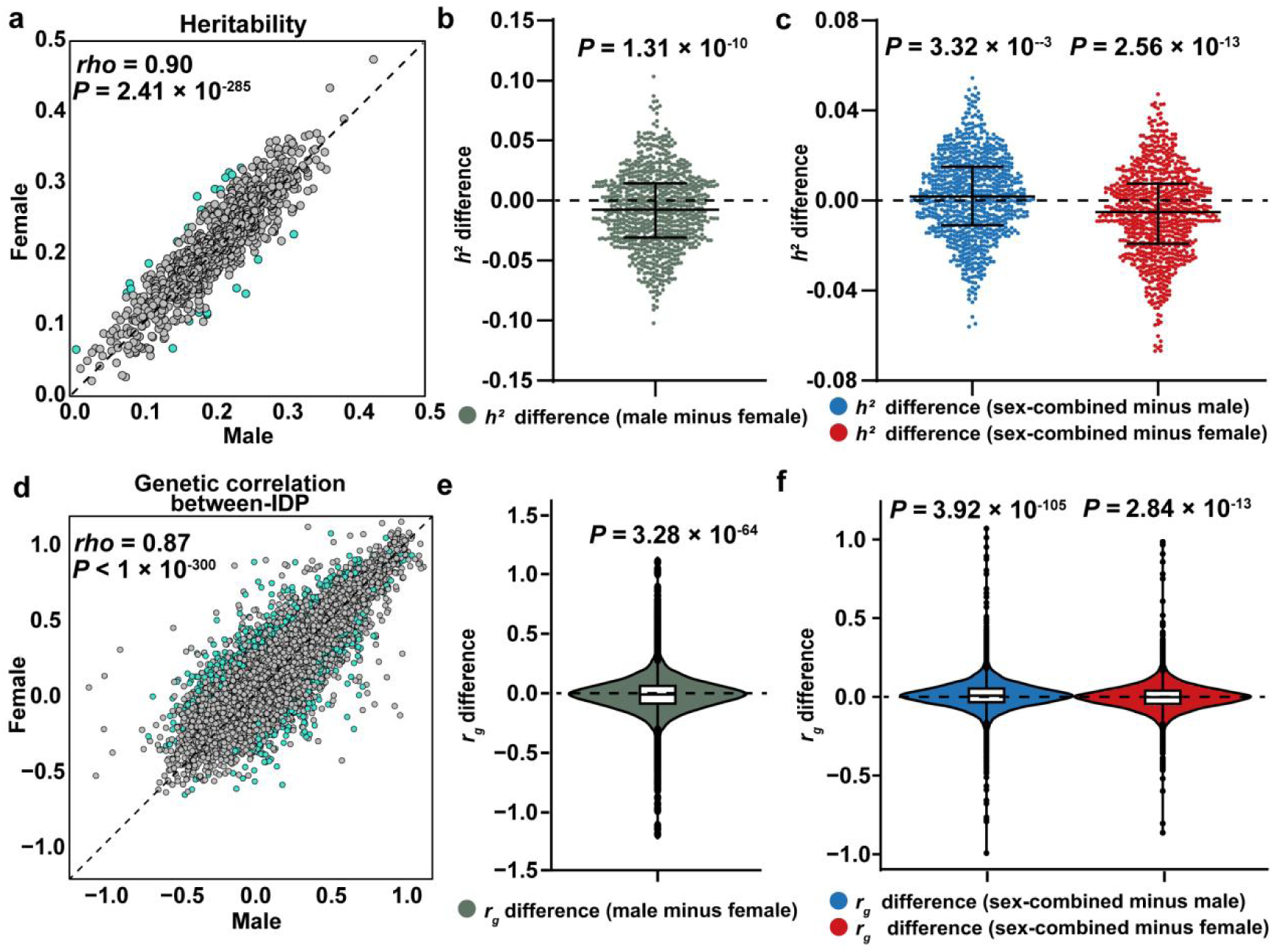
SNP-based heritability and genetic correlations. **a**, The scatter plot shows the correlation of SNP-based heritability estimates (h^2^) for 805 IDPs between males and females. The dashed line indicates the identity line, and the blue dots indicate IDPs with sex differences (two-sided P < 0.05) in heritability. **b,** Sex differences in heritability estimates for 805 IDPs. Each dot shows the heritability difference (male minus female) for each IDP. Values are statistically summarized as median difference and interquartile range (IQR). Females show higher heritability than males (paired-sample Wilcoxon signed-rank test, two-sided P = 1.31 × 10^-10^). **c,** The overall difference in heritability estimates of all IDPs between sex-stratified and sex-combined analyses. Each dot shows the heritability difference (sex-combined minus sex-stratified) of each IDP. Compared to sex-stratified analyses, sex-combined analyses overestimate heritability in males (P = 3.32 × 10^-3^) and underestimate heritability in females (P = 2.56 × 10^-13^). **d,** The scatter plot illustrates the correlation of between-IDP genetic correlations for all IDP pairs between males and females. Blue dots indicate IDPs with significant sex differences (P < 0.05) in genetic correlations. **e,** Sex differences in between-IDP genetic correlations for all IDP pairs. Inside each violin, a box plot indicates the median (central line) and IQR (box boundaries) of genetic correlation difference. Females show higher genetic correlations than males (paired-sample Wilcoxon signed-rank test, two-sided P = 3.28 × 10^-64^). **f,** The overall difference in between-IDP genetic correlations of all IDP pairs between sex-stratified and sex-combined analyses. Compared to sex-stratified analyses, sex-combined analyses overestimate genetic correlation (P = 3.92 × 10^-105^) in males and underestimate genetic correlation (P = 2.84 × 10^-13^) in females.

#### Biased heritability estimation in sex-combined sample

We performed a paired-sample Wilcoxon signed-rank test to assess the overall difference in heritability estimates of the 805 IDPs between sex-stratified and sex-combined analyses. Compared with sex-stratified analyses, sex-combined analyses overestimated heritability in males (*P* = 3.32 × 10^-3^; median *h^2^* difference = 0.0018) and underestimated heritability in females (*P* = 2.56 × 10^-13^; median *h^2^* difference = 0.0051) (Fig. 1c). The overestimated heritability in males was observed for sMRI-IDPs (*P* = 5.49 × 10^-3^) but not for dMRI-IDPs (*P* = 0.15), while the underestimated heritability in females was observed for both sMRI-IDPs (*P* = 2.07 × 10^-3^) and dMRI-IDPs (*P* = 8.75 × 10^-13^) (Extended Data Fig. 3d, e).

### Genetic correlation

Genetic correlation (*r*_g_) evaluates the shared genetic architecture between two traits measured in the same sample or between two subgroups of the same trait. We applied LDSC v.1.0.0^22^ to estimate the between-sex genetic correlation for each IDP, along with the between-IDP genetic correlation within each sex. We focused on sex differences in both kinds of genetic correlations.

#### Between-sex genetic correlation

Among the 805 IDPs, we successfully calculated the between-sex genetic correlations (ranging from 0.55 to 1.19) for 757 IDPs. We defined a sex difference as a between-sex *r*_g_ value for an IDP that is significantly lower than 1 (*P* < 0.05/805 = 6.21 × 10^-5^). Although no IDPs showed significant sex differences in between-sex genetic correlations at *P* < 6.21 × 10^-5^, 35 IDPs showed significant sex differences at *P* < 0.05 (Supplementary Table 6).

#### Between-IDP genetic correlation

We focused on 29,520 genetic correlations among IDPs for the same imaging measure using the same parcellation scheme. Each between-IDP genetic correlation was estimated in the three samples, respectively.

We assess sex differences in between-IDP genetic correlations from three aspects. First, we calculated Spearman’s rank correlation of genetic correlations between males and females, identifying strong but not perfect correlations in *r*_g_ values across all IDP pairs (*rho* = 0.87, *P* < 1 × 10^-300^; Fig. 1d), sMRI-IDP pairs (*rho* = 0.82, *P* < 1 × 10^-300^; Extended Data Fig. 4a), and dMRI-IDP pairs (*rho* = 0.90, *P* < 1 × 10^-300^; Extended Data Fig. 4b). Second, we conducted a paired-sample Wilcoxon signed-rank test to compare the overall difference in *r*_g_ values between males and females, revealing a higher genetic correlation in females than in males (*P* = 3.28 × 10^-64^; median *r_g_* difference = 0.0105; Fig.1e). This sex difference was observed for sMRI-IDPs (*P* = 4.45 × 10^-90^; median *r_g_* difference = 0.0183) but not for dMRI-IDPs (*P* = 0.98; median *r_g_* difference = 0.0005) (Extended Data Fig. 4c). Third, we performed a *Z*-test to identify sex difference in genetic correlation estimates for each IDP pair. While no differences were significant at Bonferroni-corrected *P* < 0.05/29,520 = 1.69 × 10^-6^, 1,260 genetic correlations showed significant sex differences at *P* < 0.05 (818 for sMRI-IDP pairs and 442 for dMRI-IDP pairs; Supplementary Table 7). Eight genetic correlations showed significant sex differences at *P* < 0.001, all within dMRI-IDP pairs (Extended Data Fig. 4d), such as the genetic correlation between ICVF in the left superior fronto-occipital fasciculus and in the right uncinate fasciculus (*r*_g_ = 0.36 in males, *r*_g_ = 0.68 in females, *Z* = -4.08, *P* = 4.44 × 10^-5^).

To evaluate bias in sex-combined genetic correlation estimation, we compared the overall difference in 29,520 genetic correlations estimated by sex-stratified and sex-combined analyses using the Wilcoxon signed-rank test. Compared with sex-stratified analyses, sex-combined analyses overestimated genetic correlation in males (*P* = 3.92 × 10^-105^; median *r_g_* difference = 0.0079) and underestimated genetic correlation in females (*P* = 2.84 × 10^-13^; median *r_g_* difference = 0.0024) ( Fig. 1f). For sMRI-IDP pairs, sex-combined analyses overestimated genetic correlation in males (*P* = 8.96 × 10^-70^; median *r_g_* difference = 0.0085 and underestimated genetic correlation in females (*P* = 1.69 × 10^-68^; median *r_g_* difference = 0.0101) (Extended Data Fig. 4e). For dMRI-IDP pairs, however, sex-combined analyses overestimated genetic correlation in both males (*P* = 1.59 × 10^-38^; median *r_g_* difference = 0.0069) and females (*P* = 2.89 × 10^-46^; median *r_g_* difference = 0.0073) (Extended Data Fig. 4f).

### Genome-wide associations

We conducted GWASs for 805 IDPs across 16,013,685 autosomal variants in the male, female, and sex-combined samples, respectively. The genomic control inflation factors (λ_GC_)^23^ of these GWASs ranged from 1.00 to 1.27, and their LD score regression (LDSC) intercepts^22^ ranged from 0.98 to 1.06 (Supplementary Table 4), indicating no notable population stratification. At the study-wide significance threshold (*P* < 6.21 × 10^-11^, Bonferroni corrected for 805 IDPs), we identified 2,108, 1,962, and 8,726 independent variant-trait associations from male-only, female-only, and sex-combined GWASs (Fig. 2a; Extended Data Fig. 5a; Supplementary Tables 8-10). We found the most significant association between variants at chr15 and the SA of the right precentral cortex across the three samples (male: rs5812099, *P* = 4.90 × 10^-139^; female: rs530653020, *P* = 2.82 × 10^-141^; sex-combined: rs530653020, *P* = 5.37 × 10^-278^). The two variants are linked to *THBS1*, which plays an important role in synaptogenesis^24^.

**Fig. 2.**
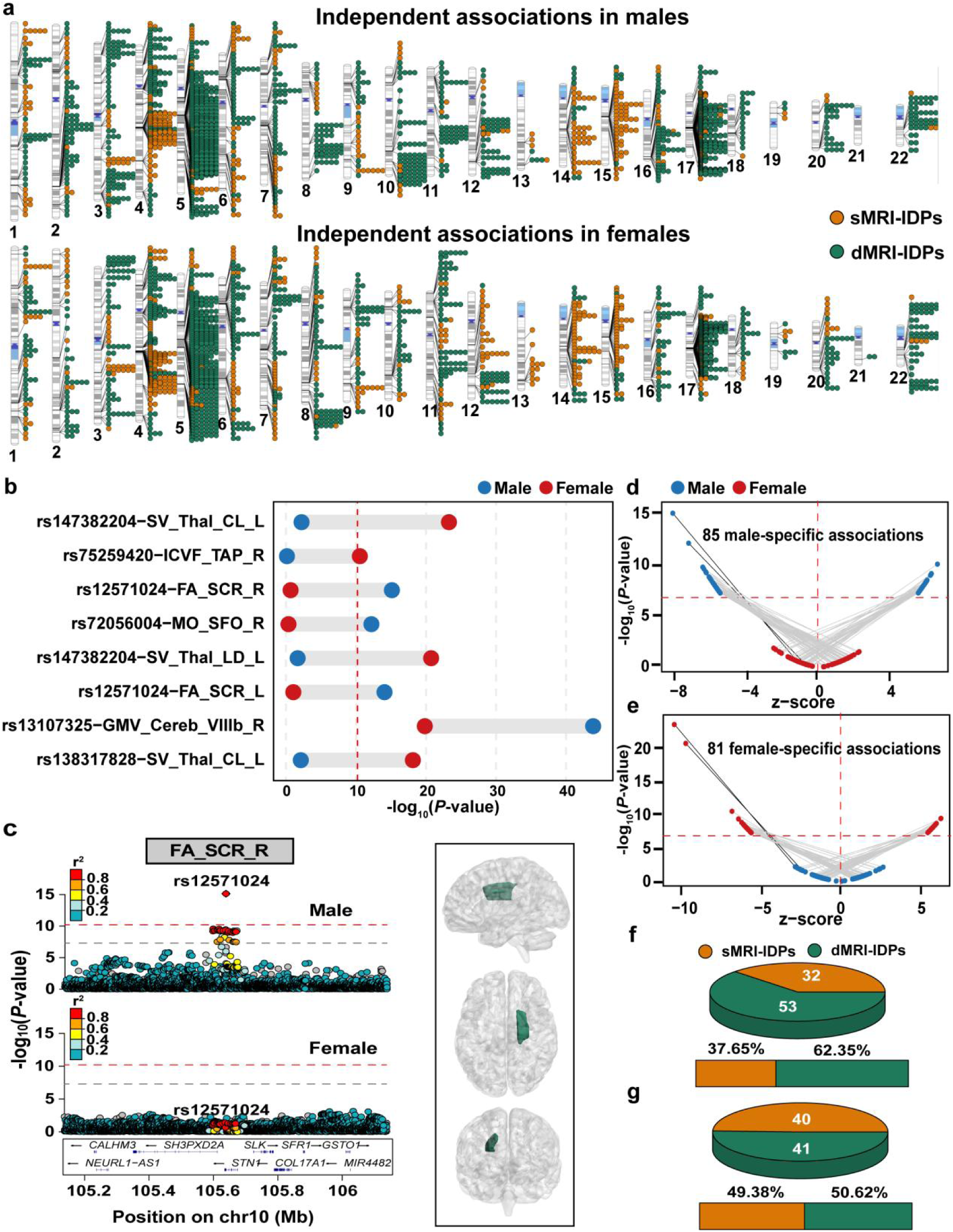
Sex differences in variant-trait associations. **a,** Ideograms depict genomic locations of independent variant–trait associations identified by male (upper panel) and female (lower panel) GWASs at P < 6.21 × 10^-11^ (Bonferroni corrected for 805 IDPs). Orange and green dots indicate associations for sMRI-IDPs and dMRI-IDPs, respectively. **b,** Independent variant–IDP associations (P < 6.21 × 10^-11^) showing significant allele-effect heterogeneity between sexes (CQ test, P < 0.05/3,265). Points indicate −log_10_(P-value) in male (blue) and female (red) GWASs; the dashed vertical line denotes the study-wide significance (P = 6.21 × 10^-11^). **c,** Locus zoom plot (left panel) shows a male-specific association between rs12571024 (chr10) and fractional anisotropy in the right superior corona radiata (right panel). **d, e,** Genetic effects (Z-scores) and GWAS significance (−log_10_(P-value)) for the 85 male-specific (**d**) and 81 female-specific (**e**) associations identified from associations significant at P < 5 × 10^-8^. Gray lines indicate associations with opposite effect directions between sexes, while black lines indicate associations with concordant effect directions. **f, g,** Pie charts show the proportions of sMRI-IDP-related and dMRI-IDP-related associations in male-specific (**f**) and female-specific (**g**) associations, respectively.

#### Sex difference in SNP effects

We aggregated all significant variant-trait associations (*P* < 6.21 × 10^-11^) identified by sex-stratified GWASs, pooling them into 3,265 unique independent variant-trait associations (Extended Data Fig. 5b; Supplementary Table 11). For each association, we performed a Cochran’s *Q* test (CQ-test)^25^ to evaluate the allele-effect heterogeneity (*P* < 0.05/3,265 = 1.53 × 10^-5^, Bonferroni corrected) between sexes, resulting in eight associations with significant sex difference (Fig. 2b; Supplementary Table 12). For example, the associations between rs12571024 (chr10) and FA in the right superior corona radiata showed heterogeneity between sexes (CQ test *P* = 6.54 × 10^-7^). The effect was significant in the male GWAS (*β* = -0.056, *P* = 7.43 × 10^-16^), but not in the female GWAS ( *β* = -0.008, *P* = 0.22) (Fig. 2c).

Given the limited sample size (n = 22,950) in sex-stratified GWASs, the study-wide significance threshold (*P* < 6.21 × 10^-11^) may overlook important associations with significant sex difference. Therefore, we used the genome-wide significance threshold (*P* < 5 × 10^-8^) to select associations, generating 10,662 unique independent variant-trait associations (Supplementary Table 13). We then performed the CQ-test to evaluate their allele-effect heterogeneity between sexes (*P* < 0.05/10,662 = 4.69 × 10^-6^, Bonferroni corrected). We identified 166 associations with significant sex differences, including 85 male-specific associations (32 for sMRI-IDPs and 53 for dMRI-IDPs; Fig. 2d, e) and 81 female-specific associations (40 for sMRI-IDPs and 41 for dMRI-IDPs; Fig. 2f, g) (Supplementary Table 14).

#### Sex-stratified versus sex-combined GWASs

We investigated if sex-stratified GWASs could offer new insights into the genetic architecture of brain IDPs compared to sex-combined GWASs. We pooled significant variant-trait associations (*P* < 6.21 × 10^-11^) identified by either sex-stratified or sex-combined GWASs into 9,796 unique independent variant-trait associations (Extended Data Fig. 5c and Supplementary Table 15). Using a sex-combined LD reference, we defined a new association when the variant was significant in male or female GWASs, but not in sex-combined GWASs, where the variant was 500 kb away from and not in LD (*r*^2^ < 0.1) with any variants in the variant-trait associations from sex-combined GWASs. Sex-stratified GWASs identified 47 new associations (Fig. 3a and Supplementary Table 16) that were not significant in sex-combined GWASs despite the sample size being doubled. These new associations involved 43 unique IDPs; most were observed for dMRI-IDPs (74.47%). For example, four genetic associations with L1 in the left posterior thalamic radiation were significant in the female GWAS but not in the male or sex-combined GWASs (Fig. 3b). These results indicate that sex-stratified GWASs could offer new insights into the genetic architecture of brain IDPs compared to the sex-combined GWASs.

**Fig. 3.**
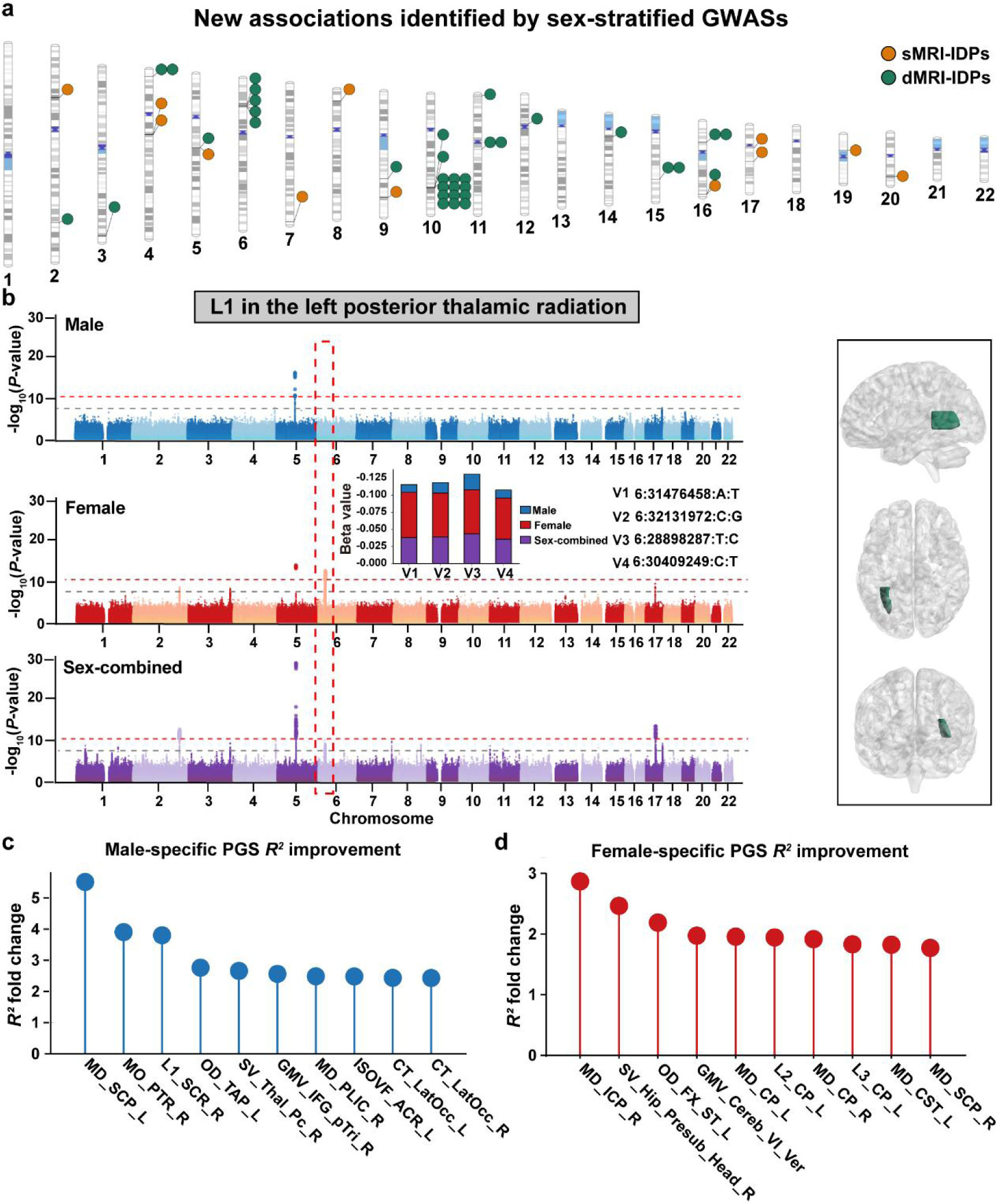
Association discovery and PGS prediction. **a,** New associations discovered by sex-stratified GWASs. Each dot denotes a new independent variant–trait association (P < 6.21 × 10^-11^) identified in sex-stratified GWASs but not (≥500 kb away and r^2^ < 0.1) in sex-combined GWASs. Orange and green points indicate sMRI-IDPs and dMRI-IDPs, respectively. **b,** Manhattan plots (left panel) show genetic associations with L1 in the left posterior thalamic radiation (right panel) that are significant in the female GWAS (middle) but not in the male (upper) or sex-combined (lower) GWAS. The x-axis denotes chromosomal position, and the y-axis indicates association strength (-log_10_P). The gray and red lines mark significance at P = 5 × 10^-8^ and P = 6.21 × 10^-11^, respectively. Insets show between-sex beta comparisons for the four significant variant-trait associations. Manhattan plots are generated as described previously^27^. **c, d,** The top 10 brain IDPs that are better predicted by the male-specific (**c**) or female-specific (**d**) PGS models compared to the sex-combined models. Fold change is calculated as the ratio of the variance explained (R²) by the sex-specific PGS model to that explained by the sex-combined PGS model. Each point represents an individual IDP, and vertical lines indicate the corresponding fold change in predictive accuracy. Please see full names for abbreviations in Supplementary Table 3.

### Polygenic score (PGS) prediction

We investigated if sex-stratified PGS models outperformed sex-combined PGS models in sex-specific phenotype prediction. To exclude bias from differences in sample sizes used for model construction, we down-sampled to randomly select 11,475 males and 11,475 females to create a sex-combined sample (n = 22,950) matching the sample size of the sex-stratified GWASs^15^. We focused on 540 IDPs with at least one significant association (*P* < 6.21 × 10^-11^) in male GWASs for male-specific prediction and 544 IDPs with at least one significant association (*P* < 6.21 × 10^-11^) in female GWASs for female-specific prediction. We utilized PRSice-2 (v.2.3.5)^26^ to construct a male-specific model from male GWAS and a sex-combined model from sex-combined GWAS for each IDP in male-specific prediction, and a female-specific model from female GWASs and a sex-combined model from sex-combined GWASs for each IDP in female-specific prediction. The target samples (n = 3,864-6,653 across IDPs) did not overlap with the base sample and comprised Caucasian participants excluded from GWAS analyses due to incomplete data for all 805 IDPs or mismatches in PSM. We used both male-specific and sex-combined models to predict IDPs for males, and both female-specific and sex-combined models to predict IDPs for females in the target sample. We calculated the variance explained (*R*^2^) by each model to evaluate predictive performance and assessed differences between models using paired-sample Wilcoxon signed-rank test.

The predictive performance (*R*^2^) of sex-specific and sex-combined PGS models is provided in Supplementary Tables 17 and 18. Although we did not identify significant differences in predictive performance between sex-specific and sex-combined models across all IDPs, sMRI-IDPs, and dMRI-IDPs (all *P* > 0.05), we found that 258 IDPs were better predicted by the male-specific model (Fig. 3c), and 276 IDPs were better predicted by the female-specific model (Fig. 3d). For example, compared with the sex-combine model, the male-specific model substantially improved prediction for mean diffusivity (MD) in the left superior cerebellar peduncle (*R^2^* = 1.33% vs. 0.24%), while the female-specific model improved prediction for MD in the right inferior cerebellar peduncle (*R^2^* = 1.75% vs. 0.61%). These findings suggest that sex-specific models may improve prediction for a subset of brain IDPs, and in some cases produce substantially higher predictive accuracy than sex-combined models.

### Statistical fine-mapping

Statistical fine-mapping can prioritize causal variants that are statistically likely to drive the observed independent variant-trait associations from GWASs. Based on the sample-specific LD reference, we conducted statistical fine mapping on independent variant-trait associations (*P* < 6.21 × 10^-11^) from each GWAS. For each association, we applied FINEMAP v1.4.1^28^ to estimate the posterior inclusion probability (PIP) for each variant within a 1000-kb locus centered on the lead variant. From these results, we created the 95% credible sets (CSs) and identified causal variants with PIP > 0.8. For each causal variant, we utilized the Ensembl Variant Effect Predictor (VEP) tool^29^ to predict its functional consequence based on genic location, and assessed its deleteriousness by calculating the combined annotation-dependent depletion (CADD) score^30^ and loss-of-function observed/expected upper bound fraction (LOEUF) value^31^.

#### Sex difference

For each of the 3,265 independent variant-trait associations from the sex-stratified GWASs, we expanded the variant into a 1Mb locus, then performed fine-mapping on that locus using the male and female GWASs, respectively. This analysis identified 276 causal variant-trait associations (PIP > 0.8) from the male GWASs and 242 from the female GWASs (Supplementary Tables 19 and 20), corresponding to 66 and 65 unique variants. We identified four missense variants (rs13107325, rs2267161, rs4935898, and rs41298373) from both male and female GWASs, and three others only in female GWASs (Supplementary Tables 21 and 22). The rs13107325 on chr10 showed a high CADD score (23.80) and a low LOEUF score (0.47); it is located in *SLC39A8*, a gene associated with the development and function of dopaminergic neurons^32^. To characterize sex differences, we defined male-dominant causal associations as those with PIP > 0.8 in males and the absolute PIP difference (ΔPIP) between sexes exceeding 0.3. The same criteria were used for defining female-dominant causal associations. We identified 133 male-dominant associations (39 for sMRI-IDPs and 94 for dMRI-IDPs) and 102 female-dominant associations (50 for sMRI-IDPs and 52 for dMRI-IDPs) (Fig. 4a, b; Extended Data Fig. 6a, b; Supplementary Tables 23 and 24). We identified 42 male-dominant associations between rs12571024 (chr10) and dMRI-IDPs. For instance, the association between rs12571024 and L3 in the left anterior corona radiata exhibited a strong male-dominant causal effect (PIP = 0.99 in males; PIP = 1.63 × 10^-6^ in females; Fig. 4c), while the association between rs150326586 (chr3) and L1 in the right cerebral peduncle showed a pronounced female-dominant effect (PIP = 3.74 × 10^-4^ in males; PIP = 1.00 in females; Fig. 4d).

**Fig. 4.**
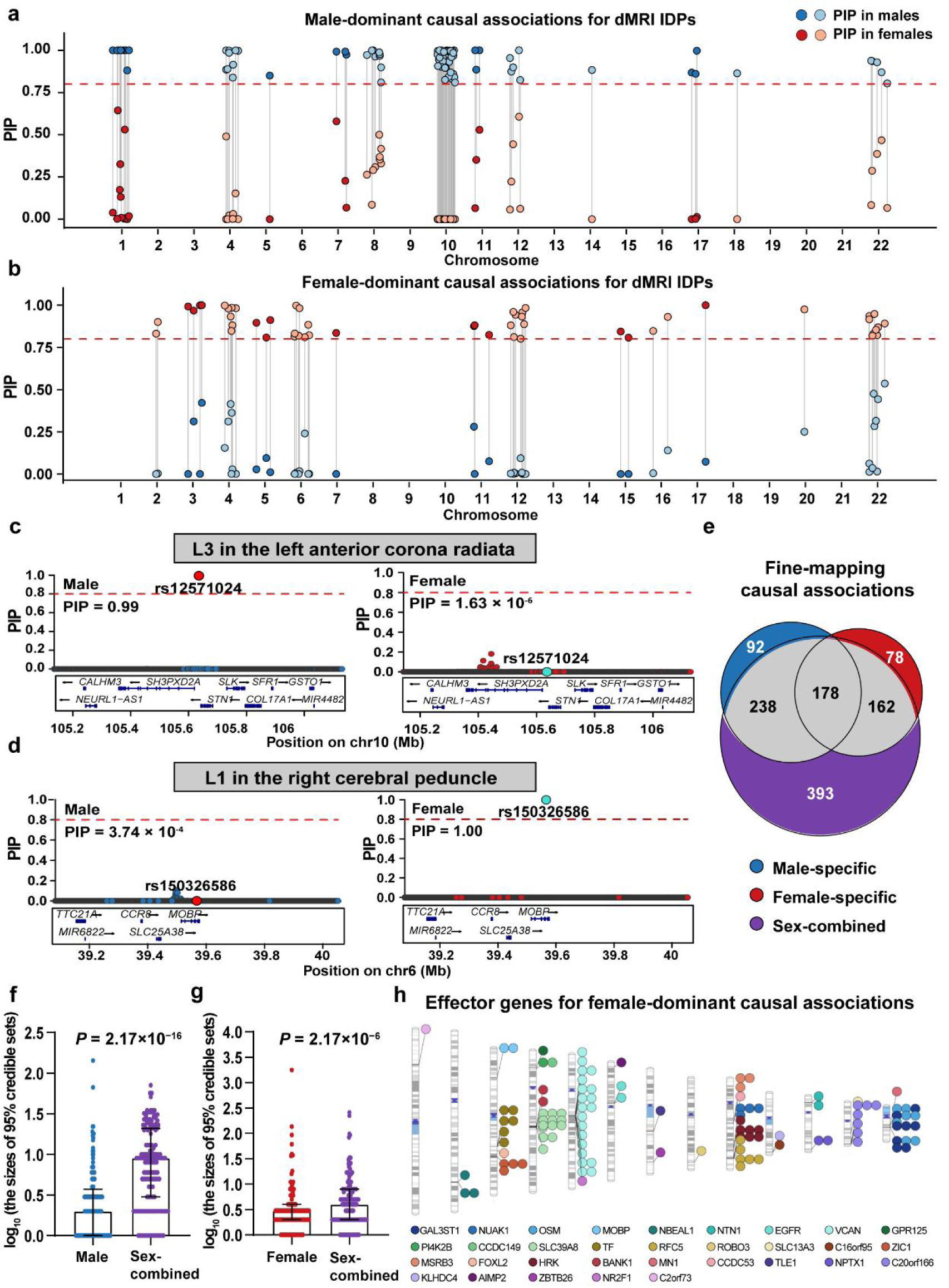
Statistical fine-mapping and gene prioritization. **a, b,** Male-dominant (**a**) and female-dominant (**b**) causal variant-trait associations for dMRI-IDPs from statistical fine-mapping. Each pair of connected points represents a variant-trait association with a PIP > 0.8 in one sex and an absolute PIP difference (ΔPIP) between sexes exceeding 0.3. Colors distinguish PIP in males (blue) and in females (red). The dashed red line indicates PIP = 0.8. **c, d,** Locus zoom plots show a male-dominant causal association between rs12571024 and L3 in the left anterior corona radiata (**c**) and a female-dominant association between rs150326586 and L1 in the right cerebral peduncle. **e,** Venn diagram illustrates the overlap and unique causal associations (PIP > 0.8) identified by male, female, and sex-combined fine-mapping analyses. **f, g,** Box plots show differences between sex-stratified and sex-combined fine-mapping in the log_10_-transformed sizes of 95% credible sets for 92 male-specific (**f**) and 78 female-specific (**g**) causal associations. P values are calculated using paired-sample Wilcoxon signed-rank test, and data are summarized as medians and IQRs. **h,** Ideogram shows the genomic locations of 32 effector genes (colors) identified from 102 female-dominant causal variant-trait associations (points), using FLAMES with a cumulative precision of >75%.

#### Sex-stratified versus sex-combined fine mapping

To determine whether sex-stratified fine-mapping can identify causal signals masked in sex-combined analyses, we pooled 9,796 independent variant-trait associations from all GWASs, and then performed fine-mapping based on the male-only, female-only, and sex-combined GWASs, respectively. We identified 508, 418, and 971 causal associations (PIP > 0.8) in the male, female, and sex-combined analyses (Supplementary Tables 25-27). Compared to the sex-combined analysis, sex-stratified analysis uncovered 170 new causal associations (Supplementary Table 28), of which 92 were found only in males (24 for sMRI-IDPs and 68 for dMRI-IDPs) and 78 were found only in females (34 for sMRI-IDPs and 44 for dMRI-IDPs) (Fig. 4e). For the sex-specific causal associations, we examined whether sex-stratified analysis could improve fine-mapping precision (the size of 95% CSs) compared with sex-combined analysis using the paired-sample Wilcoxon signed-rank test. For the 92 male-specific associations, male-specific analysis (median = 2) showed smaller CSs (*P* = 2.17 × 10^−16^; Fig. 4f) than sex-combined analysis (median = 9) (Supplementary Table 29). For the 78 female-specific associations, female-specific analysis (median = 3) also showed smaller CSs (*P* = 2.17 × 10^−6^; Fig. 4g) than sex-combined analysis (median = 4) (Supplementary Table 30). These findings indicate that sex-stratified fine-mapping may identify causal variants that cannot be identified in sex-combined samples.

#### Gene prioritization

We prioritized genes using the FLAMES framework^33^ for the 133 male-dominant and 102 female-dominant causal variant-trait associations identified in the fine-mapping analyses, corresponding to 156 and 135 locus-trait associations. Using a cumulative precision threshold of > 75%, we identified 140 locus-gene-trait pairs (36 unique genes) for male-dominant causal associations (Extended Data Fig. 6c and Supplementary Table 31) and 113 locus-gene-trait pairs (32 unique genes) for female-dominant associations (Fig. 4h and Supplementary Table 32). For example, we mapped 41 male-dominant causal associations on chr 10 to *SH3PXD2A*. This gene mediates extracellular matrix degradation^34^ and is a susceptibility gene for white matter lesions and stroke^35–38^.

#### Biological process enrichment

For the 36 effector genes from male-dominant causal associations and 32 from female-dominant causal associations, we investigated their enrichment for gene ontology (GO) terms of biological processes using WebGestalt^39^, applying a *q*_c_ < 0.05 to adjust for the Benjamini-Hochberg false positive rate (BH-FDR). We identified five terms for effector genes from male-dominant causal associations, including gliogenesis (*q*_c_ = 0.038), sensory system development (*q*_c_ = 0.038), forebrain development (*q*_c_ = 0.038), sensory organ morphogenesis (*q*_c_ = 0.043), and regulation of nervous system development (*q*_c_ = 0.043) (Extended Data Fig. 6d and Supplementary Table 33). However, no significant terms were found for effector genes from female-dominant causal associations.

### Genetic colocalization with sex hormone

A prior study^40^ provided summary statistics for both sex-stratified and sex-combined GWASs on three sex hormones: total testosterone (TT), bioavailable testosterone (BT), and sex hormone-binding globulin (SHBG), offering a unique opportunity to compare the sex-stratified and sex-combined genetic colocalizations between brain IDPs and sex hormones (Supplementary Table 34). Coloc^41^ was used to identify colocalizations for each pooled variant-IDP association (*P* < 6.21 × 10^-11^). We defined colocalization when the fourth posterior probability hypothesis (PP.H4, the probability of a shared causal variant between two traits) exceeded 0.8. We examined three types of colocalizations: (a) male-specific colocalization between male GWASs for both IDP and hormone; (b) female-specific colocalization between female GWASs for both IDP and hormone; and (c) sex-combined colocalization between sex-combined GWASs for both IDP and hormone. As both GWASs for IDP and hormone were conducted on UKB participants, we performed a sensitivity analysis to examine the influence of sample overlap between IDP-GWASs and hormone-GWASs on our results in participants with both TT and the covariates used in the original TT-GWAS^40^. Based on these data, we re-performed TT-GWAS using the same pipeline^40^ after excluding participants who were included in the IDP-GWASs. The resulting TT-GWAS summary statistics (Extended Data Fig. 7) were used for colocalization in the sensitivity analysis.

#### Sex difference

Of the 3,265 unique variant-IDP associations pooled from sex-stratified GWASs, we identified 76 male-specific and 13 female-specific colocalizations (PP.H4 > 0.8) with TT, 100 male-specific and 154 female-specific colocalizations with BT, and 103 male-specific and 58 female-specific colocalizations with SHBG (Fig. 5a and Supplementary Tables 35 and 36). We defined a male-dominant or a female-dominant colocalization as a locus showing colocalization (PP.H4 > 0.8) in males or females, with an absolute difference in PP.H4 between male and female exceeding 0.3^42^. We identified 76 male-dominant (4 for sMRI-IDPs and 72 for dMRI-IDPs) and 13 female-dominant (1 for sMRI-IDPs and 12 for dMRI-IDPs) colocalizations with TT, of which 52 (68.4%) male-dominant and 12 (92.3%) female-dominant colocalizations were replicated in the sensitivity analyses. Additionally, we identified 97 male-dominant (27 for sMRI-IDPs and 70 for dMRI-IDPs) and 126 female-dominant (90 for sMRI-IDPs and 36 for dMRI-IDPs) colocalizations for BT, and 95 male-dominant (55 for sMRI-IDPs and 40 for dMRI-IDPs) and 50 female-dominant (16 for sMRI-IDPs and 34 for dMRI-IDPs) colocalizations for SHBG (Fig. 5a-c and Supplementary Tables 37-40). For instance, a male-dominant colocalization was observed between mode of anisotropy (MO) in the posterior limb of the right internal capsule and TT (PP.H4 = 0.91 in males and PP.H4 = 0.09 in females; Fig. 5d) at the lead variant rs70600 (chr17). This intronic variant lies in *WNT3*, a key gene of the Wnt signaling pathway involved in neurodevelopment^43^. A female-dominant colocalization was observed between BT and orientation dispersion (OD) in the right external capsule (PP.H4 = 0.07 in males and PP.H4 = 0.85 in females; Fig. 5e) at rs74137403 (chr1). The shared lead variant is a well-known female-biased splicing quantitative trait locus (sQTL) linked to an exon-skipping event that causes a frameshift mutation in glioma^44^.

**Fig. 5.**
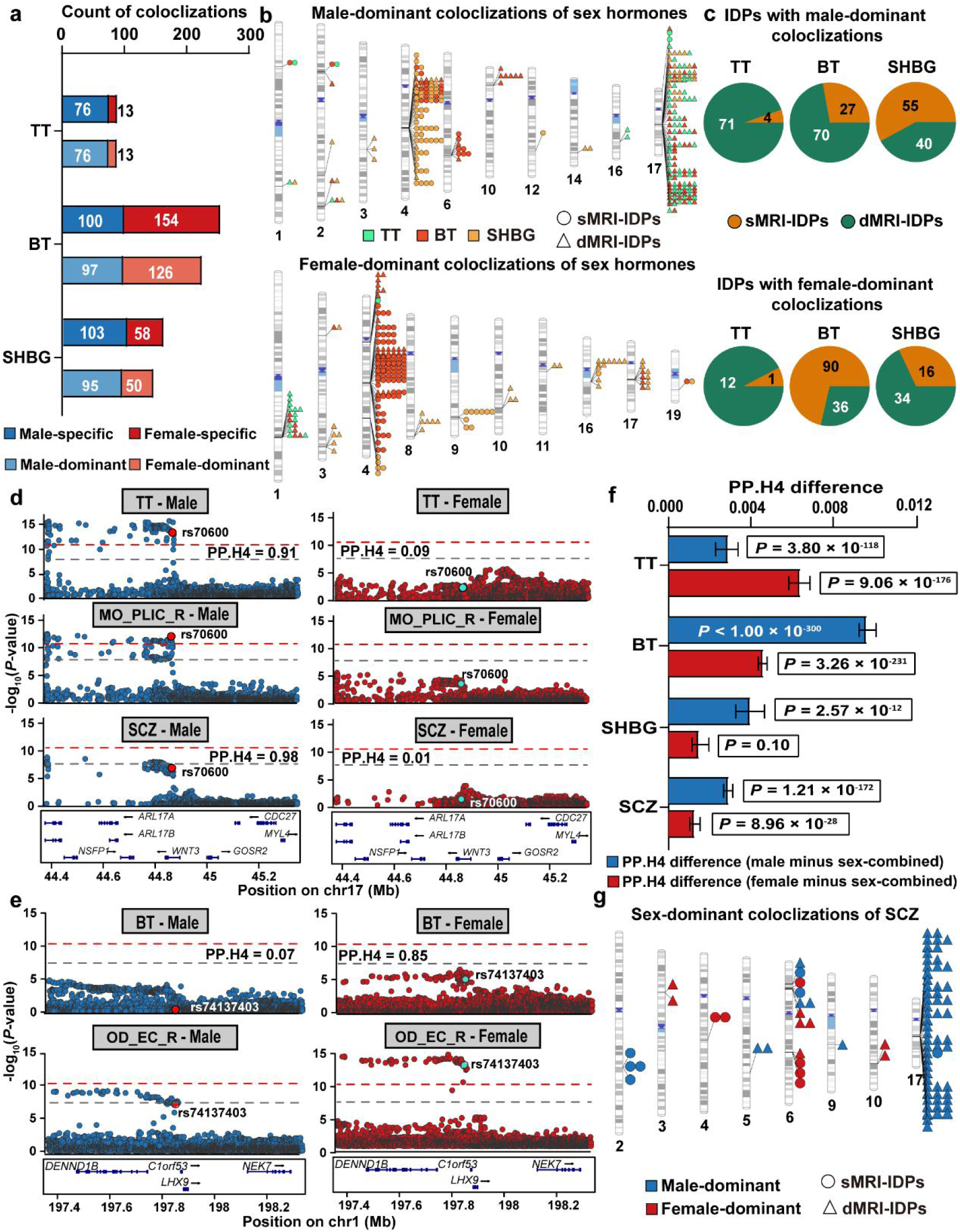
Colocalizations of brain IDPs with sex hormones and schizophrenia. **a,** Bar plots show the number of male-specific (blue), male-dominant (light blue), female-specific (red), and female-dominant (light red) colocalizations between brain IDPs and sex hormones (TT, BT, and SHBG) based on sex-stratified GWASs. **b,** Ideograms display the genomic locations of male-dominant (top panel) and female-dominant (bottom panel) colocalizations between brain IDPs (sMRI-IDPs and dMRI-IDPs) and the three sex hormones (TT, BT, SHBG). **c,** Pie charts show the percentage of sMRI-IDPs and dMRI-IDPs for male-dominant (top panel) and female-dominant (bottom panel) colocalizations with TT, BT, and SHBG, respectively. **d,** Locus zoom plots illustrate male-dominant colocalizations of mean MO in the right PLIC with TT and SCZ. **e,** Locus zoom plots illustrate a female-dominant colocalization between BT and mean OD in the right bilateral external capsule. **f,** Differences in PP.H4 (sex-stratified minus sex-combined) for colocalizations of IDPs with TT, BT, SHBG, and SCZ, with blue indicating male and red indicating female. Values are presented as median PP.H4 difference with 95% confidence interval (CI), while P values are from paired Wilcoxon signed-rank tests. Sex-stratified analyses show significantly higher PP.H4 across all traits except female-specific SHBG (P = 0.10). **g,** Ideogram shows the genomic locations of male-dominant (blue) and female-dominant (red) colocalizations between brain IDPs and SCZ. Abbreviations: BT, bioavailable testosterone; CI, confidence interval; dMRI, diffusion magnetic resonance imaging; IDP, imaging derived phenotype; MO, mode of anisotropy; OD, orientation dispersion; PLIC, posterior limb of the internal capsule; SHBG, sex hormone-binding globulin; sMRI, structural magnetic resonance imaging; TT, total testosterone; SCZ, schizophrenia.

#### Sex-stratified versus sex-combined colocalization

We also performed sex-combined colocalizations to assess whether sex-stratified analyses provide additional information compared to sex-combined analyses. Of the 9,796 unique variant-trait associations from either sex-stratified or sex-combined GWASs, we found 182, 66, and 65 colocalizations for TT, 230, 373, and 92 colocalizations for BT, and 297, 188, and 399 colocalizations for SHBG (Supplementary Tables 41-43) in the male-specific, female-specific, and sex-combined analyses. Sex-stratified analyses revealed 216 new colocalizations for TT (41 for sMRI-IDPs and 175 for dMRI-IDPs; 127 replicated in the sensitivity analysis), 558 for BT (325 for sMRI-IDPs and 233 for dMRI-IDPs) and 245 for SHBG (97 for sMRI-IDPs and148 for dMRI-IDPs) (Supplementary Tables 44-45), compared with sex-combined analysis. We used the paired-sample Wilcoxon signed-rank test to compare PP.H4 values of all candidate colocalization pairs between sex-stratified and sex-combined analyses. We found that sex-stratified analyses yielded higher PP.H4 values (all *P* < 2.6 × 10^-12^) than sex-combined analyses across all hormone categories (TT, BT, and SHBG), with the only exception of female-specific colocalizations between SHBG and all IDPs (Fig. 5f). Consistently, sex-stratified analyses also showed higher PP.H4 values for both sMRI-IDPs and dMRI-IDPs compared with sex-combined analyses (all *P* < 6.5 × 10^-12^; Extended Data Fig. 8).

### Genetic colocalization with schizophrenia

After carefully searching for all common neuropsychiatric disorders, we found that only schizophrenia had openly available summary statistics (Supplementary Table 46) for both sex-stratified and sex-combined GWASs performed in non-UKB participants^45^. Therefore, we conducted the same colocalization analyses as those performed for sex hormones.

#### Sex difference

Among the 3,265 unique variant-IDP associations from sex-stratified GWASs, we identified 246 and 178 colocalizations in male-specific and female-specific analyses, respectively (Supplementary Tables 47 and 48). Applying our definition of sex dominance (PP.H4 > 0.8 in a single sex and ΔPP.H4 > 0.3), we identified 82 male-dominant (7 for sMRI-IDPs and 75 for dMRI-IDPs) and 14 female-dominant (6 for sMRI-IDPs and 8 for dMRI-IDPs) colocalizations (Fig. 5g and Supplementary Tables 49 and 50), consistent with a higher incidence of schizophrenia in men than in women^46,47^. A locus (chr 17) with rs70600 as the lead SNP, identified as the site for a male-dominant colocalization between TT and MO in the posterior limb of right internal capsule, also exhibited male-dominant colocalization with schizophrenia (PP.H4 = 0.98 in males and PP.H4 = 0.01 in females; Fig. 5d). The variant mapped to *WNT3*, a core component of the *WNT* signal pathway, has been linked to schizophrenia^48,49^.

#### Sex-stratified versus sex-combined colocalization

We also investigated whether sex-stratified analyses can provide new information compared to sex-combined approaches. For the 9,796 variant-trait associations, we identified 672, 533, and 691 colocalizations between brain IDPs and schizophrenia in the male-specific, female-specific, and sex-combined analyses (Supplementary Tables 51-53). Crucially, sex-stratified analyses revealed 191 new colocalizations for schizophrenia (33 for sMRI-IDPs, 158 for dMRI-IDPs) compared with sex-combined analyses (Supplementary Table 54). We applied the Wilcoxon signed-rank test (*P* < 0.05) to compare PP.H4 values across all potential colocalization pairs. We found that both male-specific and female-specific analyses exhibited significantly higher colocalization probabilities compared with sex-combined analyses across all IDPs (Fig. 5f), sMRI-IDPs, and dMRI-IDPs (Extended Data Fig. 8).

## Discussion

In this study, we conducted sex-stratified GWASs on 805 brain IDPs in 22,950 males and 22,950 females matched for sample size and covariates, along with sex-combined GWASs in the combined sample. We systematically compared sex differences and sex-stratified versus sex-combined analyses across six perspectives: SNP-based heritability, genetic correlation, GWAS associations, PGS prediction, statistical fine-mapping, and colocalization. The novel findings regarding sex differences included: (1) significantly higher overall genetic correlations in females than in males; (2) 85 male-specific and 81 female-specific genome-wide significant associations; (3) 133 male-dominant and 102 female-dominant causal associations; (4) 268 male-dominant and 189 female-dominant colocalizations with sex hormones, along with 82 male-dominant and 14 female-dominant colocalizations with schizophrenia. Comparing sex-stratified with sex-combined analyses, the novel findings included: (1) sex-combined analyses overestimated heritability and genetic correlation in males and underestimated heritability and genetic correlation in females; (2) sex-stratified analyses identified 47 new GWAS associations, 170 new causal associations, 1,019 new colocalizations with sex hormones, and 191 new colocalizations with schizophrenia. Compared with a prior study^7^ reporting general concordance in genetic architectures between sexes, this study provides compelling evidence for the value of sex-stratified analyses, offering novel insights into sex-specific genetic mechanisms of brain imaging phenotypes that cannot be identified in sex-combined analyses.

Two prior studies reported significant sex differences in the genetic architecture of brain IDPs: one found two genetic associations with amygdala subregional volumes^18^, and another identified mean heritability for sMRI-IDPs, two IDPs’ genetic correlations, and two genetic associations with sMRI-IDPs^6^. Consistent with the latter study^6^, we found greater female-to-male heritability for sMRI-IDPs and extended this observation to dMRI-IDPs. However, we did not find significant differences in between-sex genetic correlations for any IDPs. Although we did not replicate sex differences in the four SNP effects on IDPs, we identified 166 sex-differing SNP effects for associations significant at *P* < 5 × 10^-8^ and eight for associations significant at *P* < 6.21 × 10^-11^. The discrepancy may stem from differences in inclusion and exclusion of sex chromosomes, sample sizes, and covariate matching across studies. Nevertheless, we consider our findings more reliable as they were derived from the largest sample size with sex subgroups matched for sample size and covariates.

Beyond these two previous studies^6,18^, we examined sex differences in the genetic architecture of brain anatomical IDPs from several new perspectives. We found higher overall between-IDP genetic correlations in females than in males; 235 sex-dominant fine-mapped causal associations; and 457 and 96 sex-dominant colocalizations with sex hormones and schizophrenia, respectively. We also mapped these sex-dominant causal associations to genes and biological processes, providing new insights into sex-specific genetic control of human brain anatomy. We mapped 41 male-dominant causal variant-trait associations with dMRI-IDPs to *SH3PXD2A*, which regulates extracellular matrix degradation^34^ and is associated with white matter lesions^35,36^, stroke^37,38^, and schwannomas^50,51^. We identified 76 male-dominant (72 for dMRI-IDPs) and 13 female-dominant (12 for dMRI-IDPs) colocalizations with total testosterone; the high male-to-female ratio is consistent with its high blood levels in men. These colocalizations were mainly found for dMRI-IDPs, suggesting a pronounced effect of testosterone on brain white matter microstructure. We found 82 male-dominant (7 for sMRI-IDPs and 75 for dMRI-IDPs) and 14 female-dominant (6 for sMRI-IDPs and 8 for dMRI-IDPs) colocalizations with schizophrenia. The high male-to-female ratio is consistent with the higher prevalence of this disorder in men than in women^46,47^, while the more male-dominant colocalizations for dMRI-IDPs align with the disconnection hypothesis of schizophrenia^52–55^.

The most important contribution of this study is the identification of novel insights from systematical comparisons between sex-stratified and sex-combined analyses. We found that sex-combined analyses overestimated heritability and genetic correlation in males and underestimated heritability and genetic correlation in females compared with sex-stratified analyses, highlighting the value of sex-stratified analyses. We also found that sex-specific PGS models may improve prediction for a subset of IDPs, although differences in predictive performance between sex-specific and sex-combined models across IDPs are modest (*P* > 0.05). While sample sizes were halved, sex-stratified analyses revealed many new associations, causal associations, and colocalizations with sex hormones and schizophrenia that could not be detected in sex-combined analyses. These results further underscore the importance of conducting sex-stratified analyses. Due to the limited availability of sex-stratified GWAS summary statistics, we cannot assess the added value of sex-stratified analyses on other traits. Therefore, we propose routinely performing sex-stratified GWASs to better understand the biology driving sex disparities in human traits and to develop personalized treatments for disorders.

Using a larger sample size and more balanced sex subgroups, we extended earlier studies by conducting several novel analyses. We not only identified more associations showing sex differences, but also discovered many sex-dominant causal associations and colocalizations. We systematically compared the advantages of sex-stratified over sex-combined analyses, highlighting the particular value of sex-stratified approaches in identifying new associations, causal associations and colocalizations. As the field moves toward precision medicine, characterizing sex-specific genetic landscapes will be key for developing targeted therapeutic strategies that account for the distinct biological mechanisms of males and females.

## Methods

### Participants

All participants were recruited by the UK Biobank (UKB)^20^ from 22 research centers across the United Kingdom between 2006 and 2010. UKB comprised demographic, genomic, environmental, neuroimaging, and behavioral data from 502,415 participants aged 37 to 73 years. The study was approved by the National Health Service Research Ethics Service (21/NW/0157), and written informed consent was obtained from each participant. We accessed the data under application number 75556. After excluding 476 participants who withdrew from UKB before August 2025 and 437,151 participants without imputed genomic data or brain magnetic resonance imaging (MRI) data, we retained 64,788 participants (30,798 males and 33,990 females).

### Genetic data

Of the 64,788 participants, 11,203 were excluded during the sample-level genetic data quality control: 1,661 with sex inconsistency; 25 with sex chromosome aneuploidy; 119 with a principal component (PC)-adjusted heterozygosity above the mean (0.1903) or a genotyping missing rate larger than 0.05; 8,573 non-Caucasian participants; and 825 due to kinship coefficient > 0.0884^56^. The remaining 53,585 participants (25,784 males and 27,801 females) were included in the following analysis. In the variant-level quality control, we applied exclusion criteria of a minor allele frequency (MAF) ≥ 0.001, an imputation quality control score (info) ≥ 0.3, and *P* ≥ 1 × 10^-7^ in Hardy-Weinberg equilibrium (HWE) to the 93,095,623 imputed variants, retaining 16,013,685 qualified autosomal variants.

### Neuroimaging data

The acquisition, preprocessing, and quality control of brain MRI data, as well as the extraction, harmonization, and normalization of brain imaging derived phenotypes (IDPs) from the UKB, have been described elsewhere^57^. We included 805 brain IDPs (Supplementary Table 3) from both structural (sMRI) and diffusion (dMRI) MRI data. From sMRI data, we included 139 IDPs (category ID 1101) to measure the gray matter volume (GMV) of 139 brain regions obtained through volume-based segmentation, 110 IDPs (category ID 191) to assess the volume of 110 subcortical substructures derived from Freesurfer subsegmentation, and 124 IDPs (category ID 196) to evaluate the cortical thickness (CT) and surface area (SA) of 62 cortical regions obtained from surface-based segmentation. From dMRI data, we included 432 IDPs (category ID 134) to evaluate nine diffusion measures of 48 white matter fiber tracts. The three diffusion tensor eigenvalues (L1, L2, and L3) reflect diffusivity along the maximum, medium, and minimum orthogonal directions; fractional anisotropy (FA) assesses white matter integrity; mean diffusivity (MD) measures the average diffusivity of three orthogonal directions; mode of anisotropy (MO) represents directional diffusion type; orientation dispersion (OD) quantifies the diversity of directional diffusion; intracellular volume fraction (ICVF) evaluates intracellular diffusion; and isotropic volume fraction (ISOVF) indicates extracellular diffusion.

Of the 53,585 participants, we retained 48,781 (23,060 males and 25,721 females; Supplementary Table 1) participants with all 805 brain IDP data. Since the skewed data distribution would violate the assumption of normality when using a linear regression model for genetic analyses, a rank-based inverse normal transformation^58^ was applied to all IDPs.

### Covariates

We controlled for age at the MRI scan (field ID 21022), age^2^, scanning sites (field ID 54), and the top 40 genetic principal components (PCs; field ID 22009) in sex-stratified analyses, while further adjusting for sex, age × sex, and age^2^ × sex in sex-combined analyses. Additionally, we adjusted for total intracranial volume (TIV, field ID 26521), mean cortical thickness (MCT, field ID 26755 and 26856), and total surface area (TSA, field ID 26721 and 26822) in the analyses of GMV-IDPs, CT-IDPs, and SA-IDPs, respectively. Rank-based inverse normal transformation^58^ was performed to normalize all continuous covariates before the statistical analyses.

### Propensity score matching

Using the 48,781 participants (23,060 males and 25,721 females) with qualified genetic, IDP, and covariate data, we established male and female subgroups that were matched in sample size and covariates (age, age^2^, scanning sites, and the top 40 genetic PCs) by performing propensity score matching (PSM)^21^ using the MatchI v4.7.0 R package^59^. We conducted 1:1 matching using a greedy nearest-neighbor algorithm (caliper width = 0.2 without replacement). PSM generated 22,950 pairs of males and females. The two subgroups showed balance in these covariates, indicated by an absolute standardized mean difference (SMD) below 0.1 (Supplementary Table 2). The sex-combined sample (n = 45,900) consisted of all matched males (n = 22,950) and females (n = 22,950).

### Heritability

Based on single nucleotide polymorphisms (SNPs) on the autosomes, we used LDSC v.1.0.0^22^ to estimate the SNP-based heritability (*h^2^*) and its standard error (SE) for each brain IDP in the three samples, respectively. We applied the paired-sample Wilcoxon signed-rank test to compare heritability estimates of 805 brain IDPs between samples. Additionally, we calculated Spearman’s rank correlation for the heritability estimates of these traits between samples (*P* < 0.05). For each brain IDP, we further compared the difference in SNP-based heritability between males and females by calculating a *Z* score based on the established formula (1)^6^:

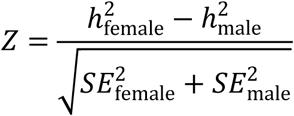

### Genetic correlation

We also used LDSC v.1.0.0^22^ to estimate the between-sex genetic correlation for each IDP, along with the between-IDP genetic correlation in each sample. We focused solely on the 29,520 between-IDP genetic correlations among IDPs for the same imaging measure and from the same parcellation. For each IDP, we identified sex difference in genetic correlation by testing whether between-sex genetic correlation was statistically significant different from *r*_g_ = 1 using the following formula (2)^19^:

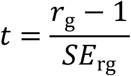

We used the paired-sample Wilcoxon signed-rank test to compare between-IDP genetic correlations between samples, and calculated their Spearman’s rank correlation between samples (*P* < 0.05). For each between-IDP genetic correlation, we compared its sex difference by calculating a *Z* score based on the established formula

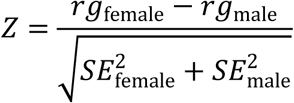

### Genome-wide association study (GWAS)

We used the imputed genotype data from the 45,900 participants (22,950 males and 22,950 females) to construct sex-combined, male-specific, and female-specific linkage disequilibrium (LD) references. For each sample, we conducted GWASs on 805 brain IDPs across 16,013,685 autosomal variants using an additive model in BGENIE v1.3^60^ (https://jmarchini.org/bgenie), while controlling for the covariates. We reported study-wide significant associations (*P* < 6.21 × 10^-11^, Bonferroni corrected for 805 IDPs). LD score regression (LDSC v.1.0.0; https://github.com/bulik/ldsc)^22^ was used to calculate the genomic control inflation factor (λ_GC_)^23^ and the LDSC intercept for each GWAS, using the sample-specific LD reference. λ_GC_ was used to evaluate the bias of the GWAS, while the LDSC intercept can differentiate between inflation caused by polygenicity and that resulting from bias (population stratification).

### Definition of independent variant-trait associations

Using a sample-specific LD reference, we performed PLINK clumping^61^ to identify independent variant-trait associations from each GWAS: (a) all variants in significant variant-trait associations were included to create a list of candidate variants; (b) of these, the most significant variant was defined as the first lead variant, and other variants within 500 kb from and in LD (*r*^2^ > 0.1) with the lead variant were clumped; (c) the remaining variants formed a new list of candidate variants, and step (b) was repeated; and (d) the iterative process continued until the list was empty. These lead variants were defined as the independent variants for the trait, and their associations were defined as the independent variant-trait associations.

### Pooling variant-trait associations

We created two sets of variant-trait associations for post-GWAS analyses using the sex-combined LD reference. For each IDP, we aggregated significant variants from any of the two sex-stratified GWASs, and then reperformed PLINK clumping^61^ to identify independent variants for the IDP. All retained independent variant-trait associations were used to investigate sex differences. We further pooled variant-trait associations from both sex-stratified and sex-combined GWASs to compare the applications of the two GWAS strategies.

### Allele-effect heterogeneity between sexes

For the pooled variant-trait associations (*P* < 5 × 10^-8^ or *P* < 6.21 × 10^-11^) from sex-stratified GWASs, we utilized Cochran’s Q test^25^ to assess allele-effect heterogeneity between sexes. Based on the Cochran’s Q test, we defined associations with *P_c_* < 0.05 (Bonferroni corrected) as the variant-trait associations with significant sex difference.

### Definition of new genetic associations

Based on the pooled variant-trait associations from sex-stratified and sex-combined GWASs, we examined whether sex-stratified GWASs could uncover new associations compared to sex-combined GWASs at *P* < 6.21 × 10^-11^. We defined a new variant-trait association when it was significant in sex-stratified GWASs but not in sex-combined GWASs, provided that the variant was 500-kb away from and not in LD (*r*^2^ < 0.1) with any significant variants of the same trait in the sex-combined GWASs.

### Polygenic score (PGS) prediction

From the 22,950 male and 22,950 female participants used in the sex-stratified GWASs, we randomly selected 11,475 males and 11,475 females to create a sex-combined cohort (*n* = 22,950) with a sample size equivalent to the sex-stratified GWASs^62^. We defined the male (*n* = 22,950), female (*n* = 22,950), and sex-combined (*n* = 22,950) participants as the base samples, while designating the 2,834 males and 4,851 females who were excluded due to incomplete data for IDPs or mismatches in PSM as two target samples. We focused solely on the IDPs that had at least one significant association in male-only GWASs for male-specific PGS prediction and on those that had at least one significant association in female-only GWASs for female-specific PGS prediction. For each IDP, the base sample was used to estimate the effect sizes of genetic variants, which were applied to construct the PGS model. We used PRSice-2 (v.2.3.5)^26^ to construct two PGS models for male-specific prediction (one based on male GWASs and another based on sex-combined GWASs) and two PGS models for female-specific prediction (one based on female GWASs and another based on sex-combined GWASs). The 2,834 males were used as the target dataset to evaluate male-specific predictions, and the 4,851 females were used as the target dataset to evaluate female-specific predictions. The performance of each PGS model was assessed by calculating the variance (*R*^2^) of the trait explained by PGS, adjusting for the covariates. For each type of sex-specific predictions, we used the paired-sample Wilcoxon signed-rank test to compare the *R*^2^ values of IDPs derived from sex-matched and sex-combined PGS models.

### Statistical fine-mapping

For each independent variant-trait association, we conducted statistical fine mapping using FINEMAP v1.4.1 (http://www.christianbenner.com/) with a shotgun stochastic search algorithm^28^, based on sample-specific LD reference and GWAS. The maximum number of causal variants was set to one. For each independent variant-trait association, we created a locus (1000-kb centered at the lead variant) and calculated the posterior probabilities (PPs) of the one causal variant model. The posterior inclusion probability (PIP) of each variant was equal to the PP of this single causal model. We also estimated the 95% credible sets of the locus based on PPs of SNPs within the locus. Specifically, the PPs of the SNPs within the locus were ordered from the largest to the smallest, and then we accumulated the PPs from the largest one until the joint probability ≥ 95%, and the list of SNPs formed the 95% credible set. PIP > 0.8 was used to identify causal variants for each brain IDP.

We identified causal variant-trait associations (PIP > 0.8) from the pooled variant-trait associations derived from sex-stratified GWASs. A male-dominant causal variant-trait association was defined as PIP > 0.8 in males and the absolute difference in PIP (ΔPIP) between sexes exceeding 0.3^42^. These criteria were also used to define female-dominant causal association. We also performed fine mapping for the pooled variant-trait associations from either sex-stratified or sex-combined GWASs. We conducted the paired-sample Wilcoxon signed-rank test (*P* < 0.05) to determine whether sex-stratified fine mapping reduce the 95% credible sets compared with the sex-combined analysis for these new causal associations.

### Functional annotations

For causal variants (PIP > 0.8) from sex-stratified GWASs, the Ensembl Variant Effect Predictor (VEP) tool^29^ (http://www.ensembl.org/info/docs/tools/vep/index.html) was used for functional annotations. We annotated each variant based on genic location and functional consequences, combined annotation-dependent depletion (CADD), and loss-of-function mutations. The CADD score predicts the deleteriousness of each variant by estimating its effect on protein structure and function; CADD > 12.37 was considered pathogenic^30^. The loss-of-function observed/expected upper bound fraction (LOEUF) assesses a gene’s sensitivity to loss of function^31^.

### Gene prioritization and pathway enrichment

We prioritized genes using the fine-mapped locus assessment model of effector genes (FLAMES)^33^, which is based on the statistical fine-mapping results and pathway-naive Polygenic Priority Score (PoPS) results from each set of GWASs. FLAMES integrates SNP-to-gene evidence with convergence-based evidence to produce a single XGBoost score for each gene. The prioritized genes were defined by cumulative precision > 75%. For the unique genes from male-dominant or female-dominant causal associations, we used WebGestalt^39^ to identify significant enrichment of these genes for gene ontology (GO) biological processes (*q*_c_ < 0.05, Benjamini-Hochberg false positive rate corrected).

### Genetic colocalizations with sex hormones

Since sex hormones influence the brain in a sex-specific manner^40^, we sought to identify male-dominant and female-dominant colocalizations between IDPs and sex hormones using coloc v5.1.0 (https://chr1swallace.github.io/coloc/index.html)^41^, and to evaluate whether sex-stratified analyses provide additional colocalization information compared with sex-combined analyses. In these analyses, we obtained both sex-stratified and sex-combined GWAS summary statistics for brain IDPs from our study, and those for total testosterone (TT), bioavailable testosterone (BT), and sex hormone-binding globulin (SHBG) from a previous study^40^. GWASs for hormones were also performed on UKB participants: 194,453 males, 230,454 females, and 425,097 sex-combined participants for TT; 178,782 males, 188,507 females, and 382,988 sex-combined participants for BT; and 180,726 males, 189,473 females, and 370,125 sex-combined participants for SHBG (Supplementary Table 34). To assess the influence of sample overlap between GWASs of brain IDPs and sex hormones, we conducted a sensitivity analysis on TT for which raw data were available. Specifically, we re-ran TT-GWASs using the same analytical pipeline and covariates as in the original study^40^, in 155,469 males, 184,078 females, and 339,547 sex-combined participants, after excluding individuals who were included in the GWASs of brain IDPs. Colocalization analyses were then repeated using the resulting summary statistics.

For variant-trait associations (*P* < 6.21 × 10^-11^) pooled from sex-stratified GWASs, we performed male-specific colocalizations using male GWASs for both IDPs and sex hormones, and female-specific colocalizations using female GWASs for both IDPs and sex hormones. Each colocalization region was defined as a 1000 kb window centered on the lead variant from the IDP-GWAS association. Using the default priors (*P*_1_ = 1 × 10^-4^, *P*_2_ = 1 × 10^-4^ and *P*_12_ = 1 × 10^-5^), the fourth posterior probability hypothesis (PP.H4, the probability of a shared causal variant between the two traits) was computed using the approximate Bayes factor^41^. We considered colocalization when PP.H4 > 0.8. We defined male-dominant or female-dominant colocalization using two criteria^42^: PP.H4 > 0.8 in either males or females, and the absolute difference in PP.H4 (ΔPP.H4) between sexes exceeding 0.3.

For the pooled variant-trait associations from either sex-stratified or sex-combined GWASs, we performed male-specific colocalizations, female-specific colocalizations, and sex-combined colocalizations between brain IDPs and sex hormones. We used the paired-sample Wilcoxon signed-rank test to examine the differences in PP.H4 values between sex-stratified and sex-combined analyses. We defined new colocalizations (PP.H4 > 0.8) as those identified in sex-stratified but not in sex-combined analyses.

### Genetic colocalizations with schizophrenia

We also investigated genetic colocalizations between brain IDPs and schizophrenia. For schizophrenia, both sex-stratified and sex-combined GWAS summary statistics were available, and none of these samples overlapped with those used in the brain IDP-GWAS analyses (Supplementary Table 46). We conducted the same analyses using the identical methods employed in the colocalization analyses for sex hormones, except for not performing the sensitivity analysis.

## Acknowledgements

This work was supported by the National Natural Science Foundation of China to C.Y and J.X (Grant No. 82430063 and 82371924), National Key Project of Inter-governmental International Scientific and Technological Innovation Cooperation to J.X (Grant No. 2023YFE0199700), the Natural Science Foundation of Tianjin to J.X (Grant No. 25JCZDJC00640), the Tianjin Young Talents in Science and Technology for J.X (Grant No. QN20230336), supported by the Non-profit Central Research Institute Fund of Chinese Academy of Medical Sciences to J.X (Grant No. 2024-JKCS-18), the Tianjin Science and Technology Commission Major Special Project in Public Health Science and Technology to J.X (Grant No. 24ZXGQSY00050) and the Tianjin Medical University Clinical Talent Training 123 Climbing Plan to J.X.

## Author contributions

Chunshui Yu and Nannan Zhang designed the study and wrote the manuscript. Nannan Zhang, Shaoying Wang, Jilian Fu and Yuan Ji analyzed the data. Chunshui Yu, Jiayuan Xu and Wen Qin supervised this work. Nannan Zhang, Shaoying Wang, Jilian Fu, Yuan Ji, Nana Liu, Qian Qian, Hui Xue, Hao Ding, Meng Liang, Wen Qin, Jiayuan Xu and Chunshui Yu acquired the data. All authors critically reviewed the manuscript.

## Data availability

All sex-stratified and sex-combined GWAS summary statistics of brain IDPs in this study are available. Interactive Manhattan plots and related data resources for this study are available at https://ssga-brain.manus.space.

## Code availability

All codes used to generate results are publicly available.

## Extended Data Figures

**Extended Data Fig. 1.**
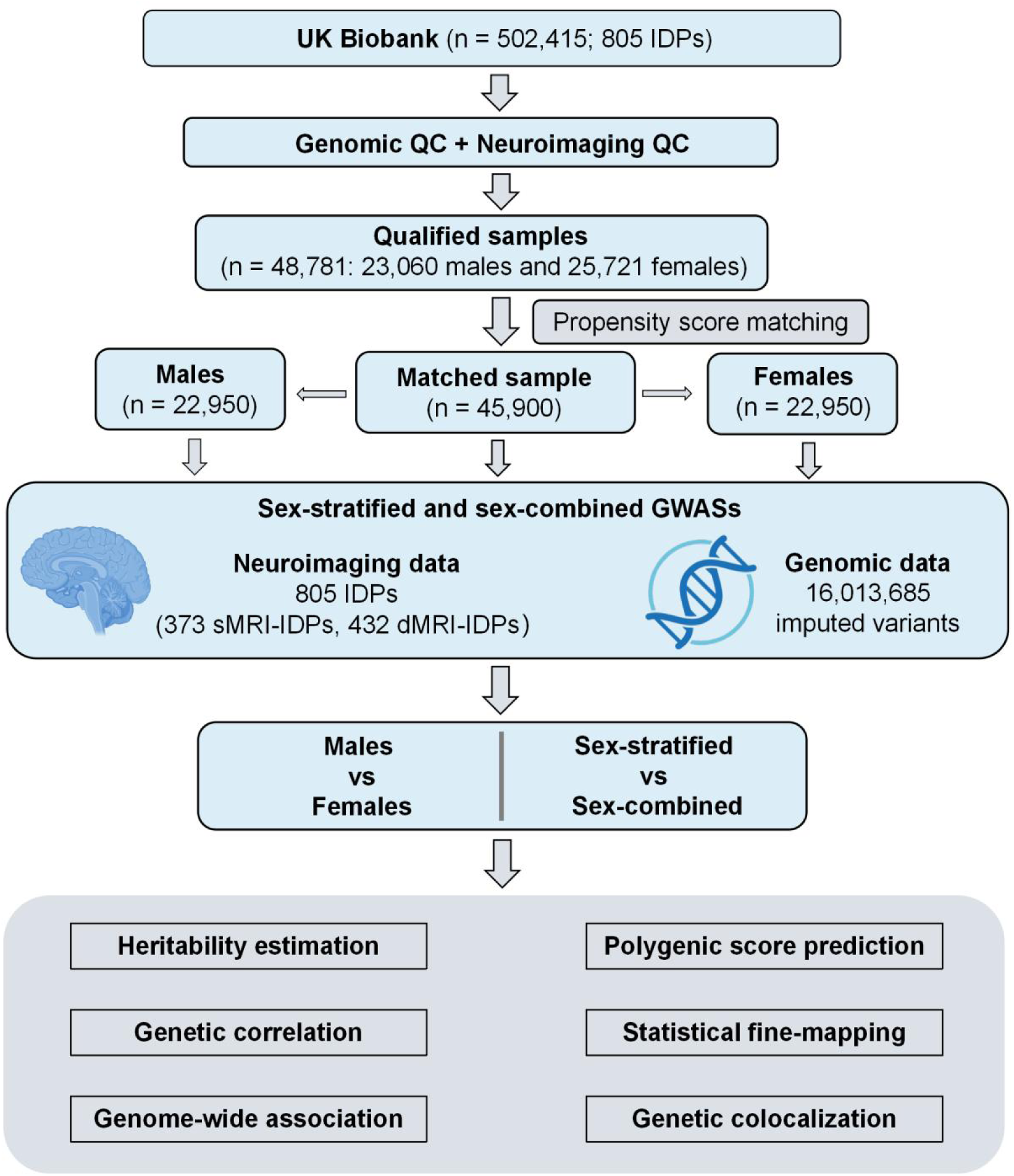
Overview of the study design. This study uses data from the UK Biobank. Following genomic and neuroimaging quality control (QC), propensity score matching (PSM) is applied to generate male and female subgroups matched for sample size (n = 22,950) and covariates; the two subgroups are then combined into a sex-combined sample (n = 45,900). We conduct both sex-stratified and sex-combined genome-wide association studies (GWASs) on 805 brain imaging-derived phenotypes (IDPs), comprising 373 for structural MRI (sMRI) and 432 for diffusion MRI (dMRI), using 16,013,685 imputed autosomal variants. We compare differences between sexes and/or between sex-stratified and sex-combined genetic analyses across six aspects: heritability estimation, genetic correlation, genome-wide association, polygenic score prediction, statistical fine-mapping, and genetic colocalization.

**Extended Data Fig. 2.**
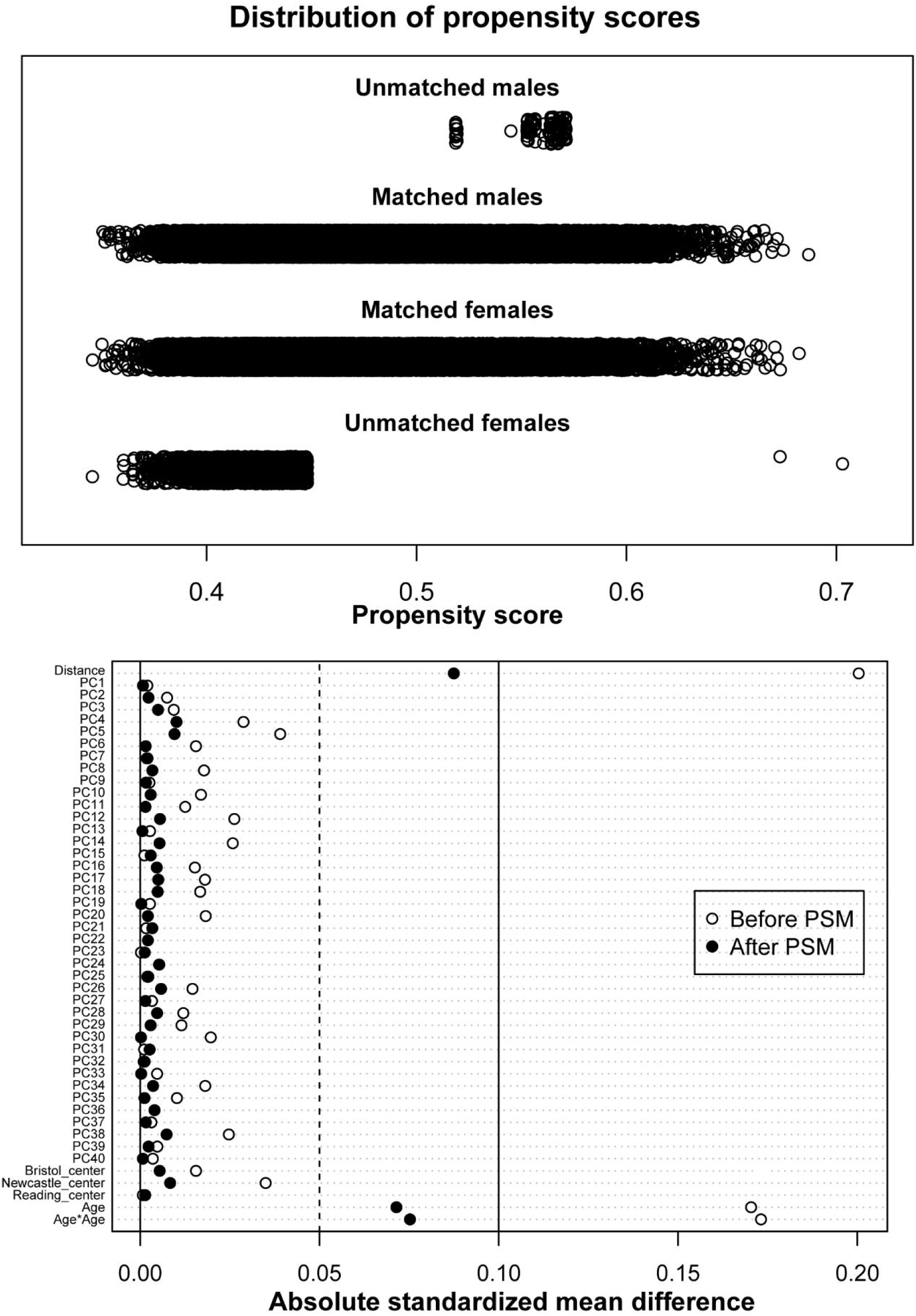
Propensity score matching. The top panel demonstrates the distribution of propensity scores before and after matching for males and females. Males and females are one-to-one matched using nearest neighbor matching based on 45 covariates, comprising age, age², three scanning sites, and the top 40 genetic principal components. The bottom panel demonstrates the absolute standardized mean differences (ASMDs) of the 45 covariates between males and females before (open circles) and after (black dots) matching. After matching, 22,950 male-female pairs are retained, with ASMDs reduced to < 0.1 for all covariates.

**Extended Data Fig. 3.**
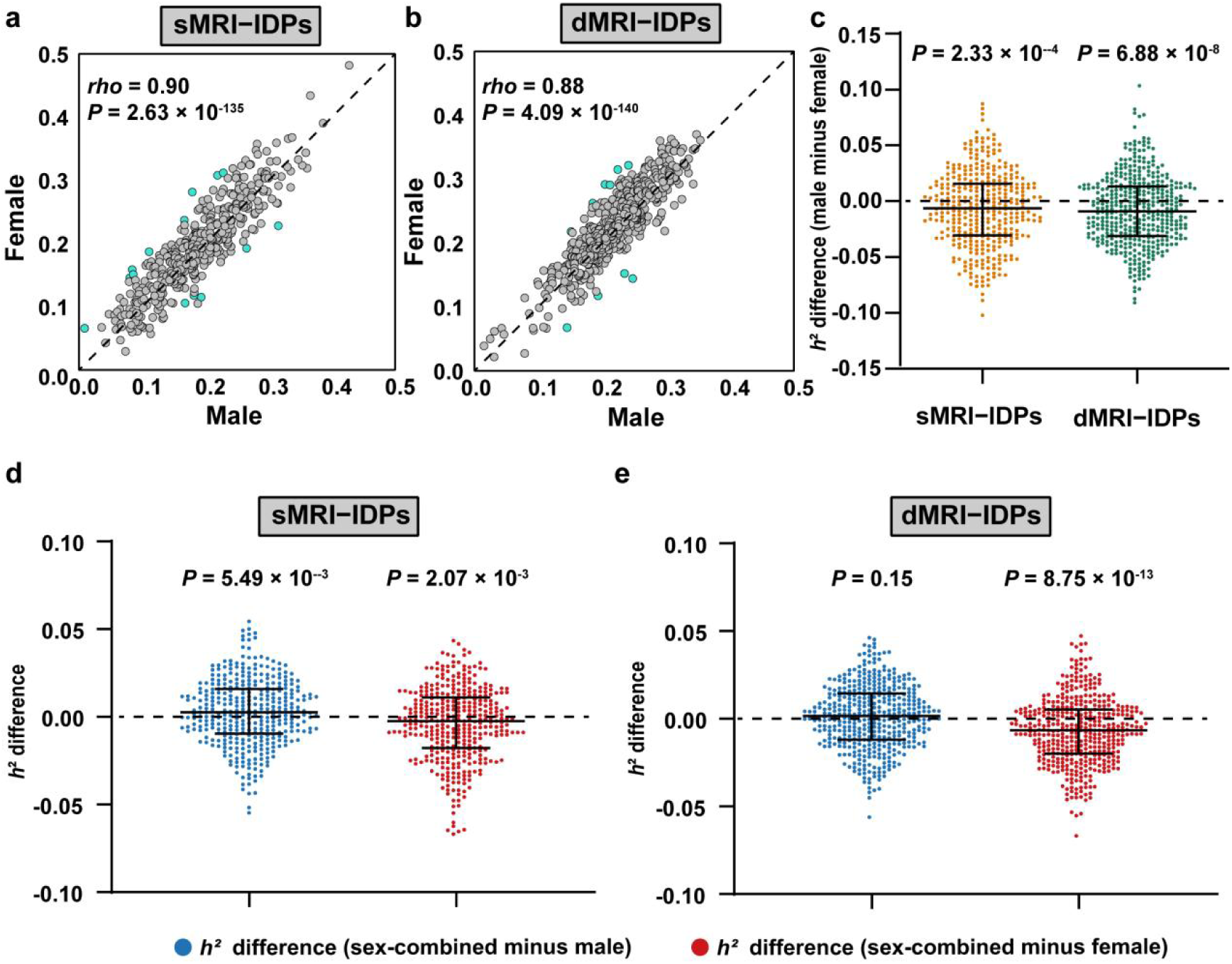
SNP-based heritability differences of brain IDPs between sexes and between sex-stratified and sex-combined analyses. **a, b,** Scatter plots show correlations of SNP-based heritability estimates between males and females for sMRI-IDPs (**a**) and dMRI-IDPs (**b**). The dashed line indicates the identity line, and the blue dots represent IDPs with sex differences (*P* < 0.05) in heritability. **c,** Sex differences (Wilcoxon signed-rank test: *P* < 0.05) in heritability for sMRI-IDPs and dMRI-IDPs. Each dot represents the heritability difference (male minus female) for an IDP. Values are summarized as median difference and interquartile range (IQR). **d, e,** SNP-based heritability differences between sex-stratified and sex-combined analyses for sMRI-IDPs (**d**) and dMRI-IDPs (**e**). Each dot represents the heritability difference (sex-combined minus sex-specific) for an IDP.

**Extended Data Fig. 4.**
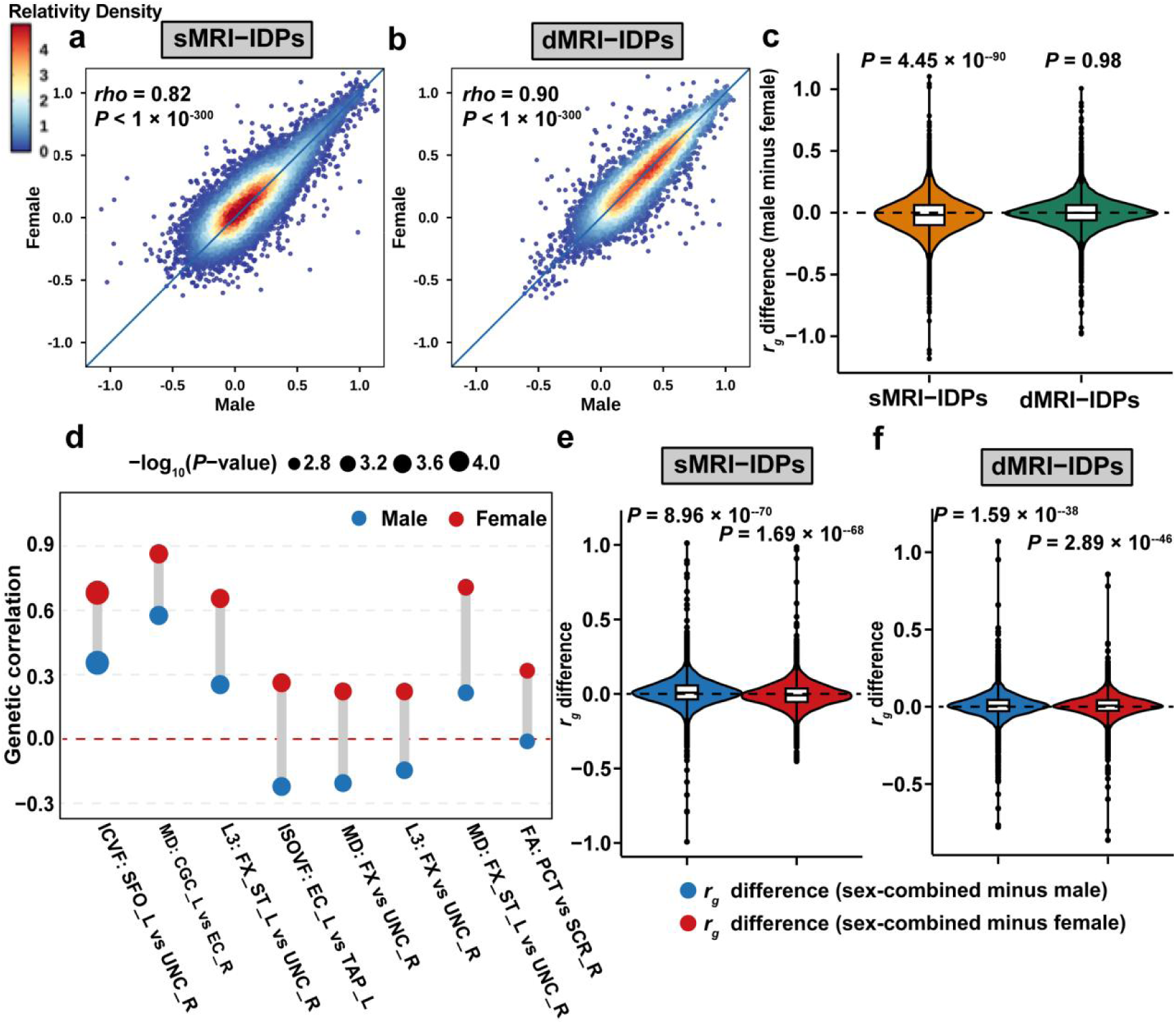
Comparisons of between-IDP genetic correlations between sexes and between sex-stratified and sex-combined analyses. **a, b,** Scatter density plots show Spearman’s correlations of between-IDP genetic correlations (r_g_) between males and females for sMRI-IDP (**a**) and dMRI-IDP (**b**) pairs. The color indicates the relative density of data points, with warmer colors indicating higher density. The blue line denotes the identity line. **c,** Violin plots demonstrate the overall sex difference in between-IDP genetic correlations for sMRI-IDP and dMRI-IDP pairs. Inside each violin, a box plot indicates the median (central line) and IQR (box boundaries) of genetic correlation difference. *P*-values are derived from the paired-sample Wilcoxon signed-rank test. **d,** Eight IDP pairs with sex differences in between-IDP genetic correlation at *P* < 0.001. Points represent r_g_ estimates in males (blue) and females (red), with point size proportional to −log_10_(*P*-value). See full names for abbreviations in Supplementary Table 3. **e, f,** The differences in between-IDP genetic correlations of sMRI-IDP (**e**) and dMRI-IDP (**f**) pairs between sex-stratified and sex-combined analyses.

**Extended Data Fig. 5.**
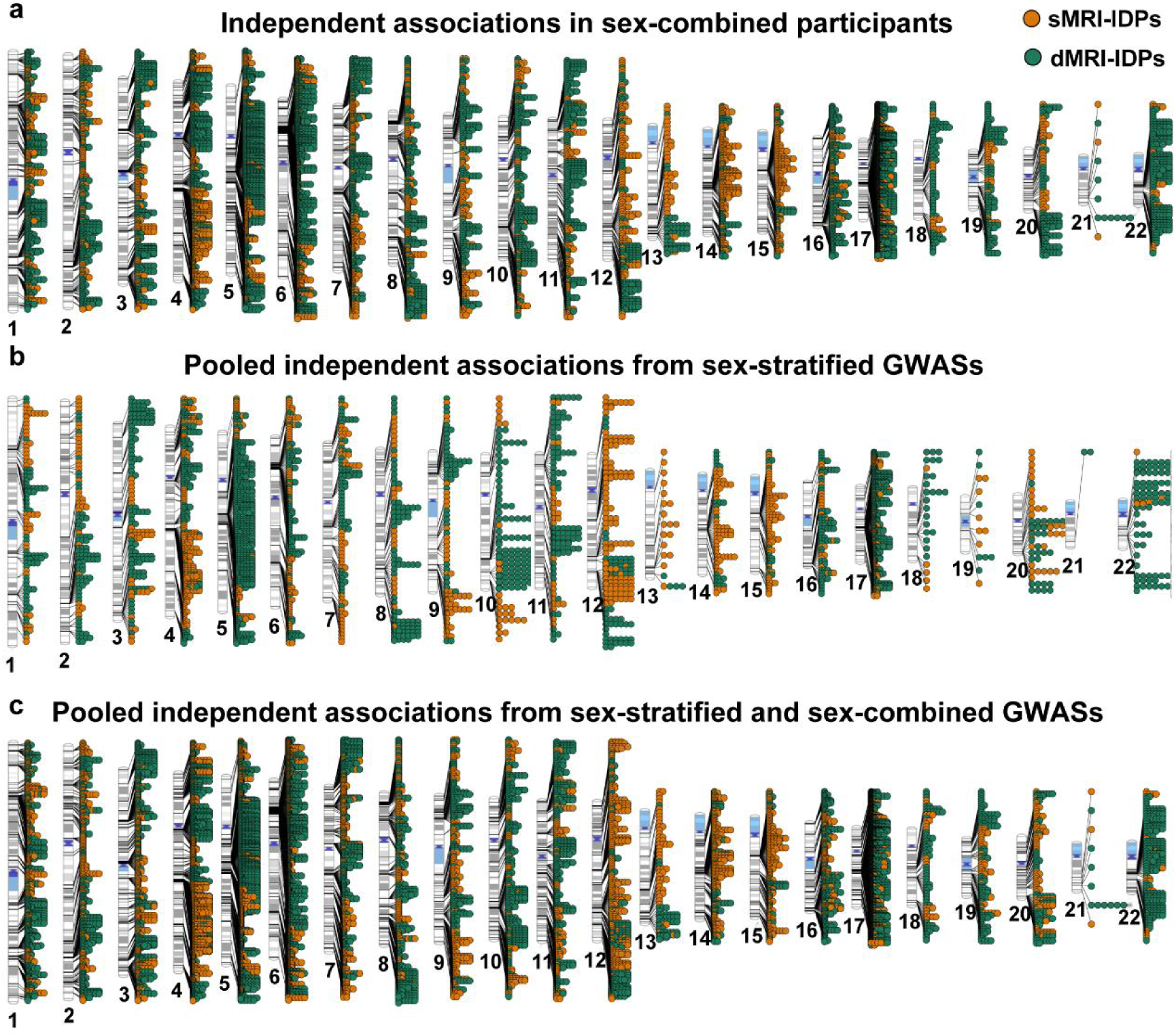
Independent variant-trait associations for brain IDPs. **a-c,** Ideograms show the genomic locations of 8,726 independent variant-trait associations identified in sex-combined GWASs (**a**), 3,265 pooled from sex-stratified GWASs (**b**), and 9,796 pooled from sex-stratified and sex-combined GWASs (**c**). Orange points denote associations with sMRI-IDPs and green points denote associations with dMRI-IDPs. The x-axis represents chromosomal position, and each dot represents a significant variant-trait association (*P* < 6.21 × 10^−11^).

**Extended Data Fig. 6.**
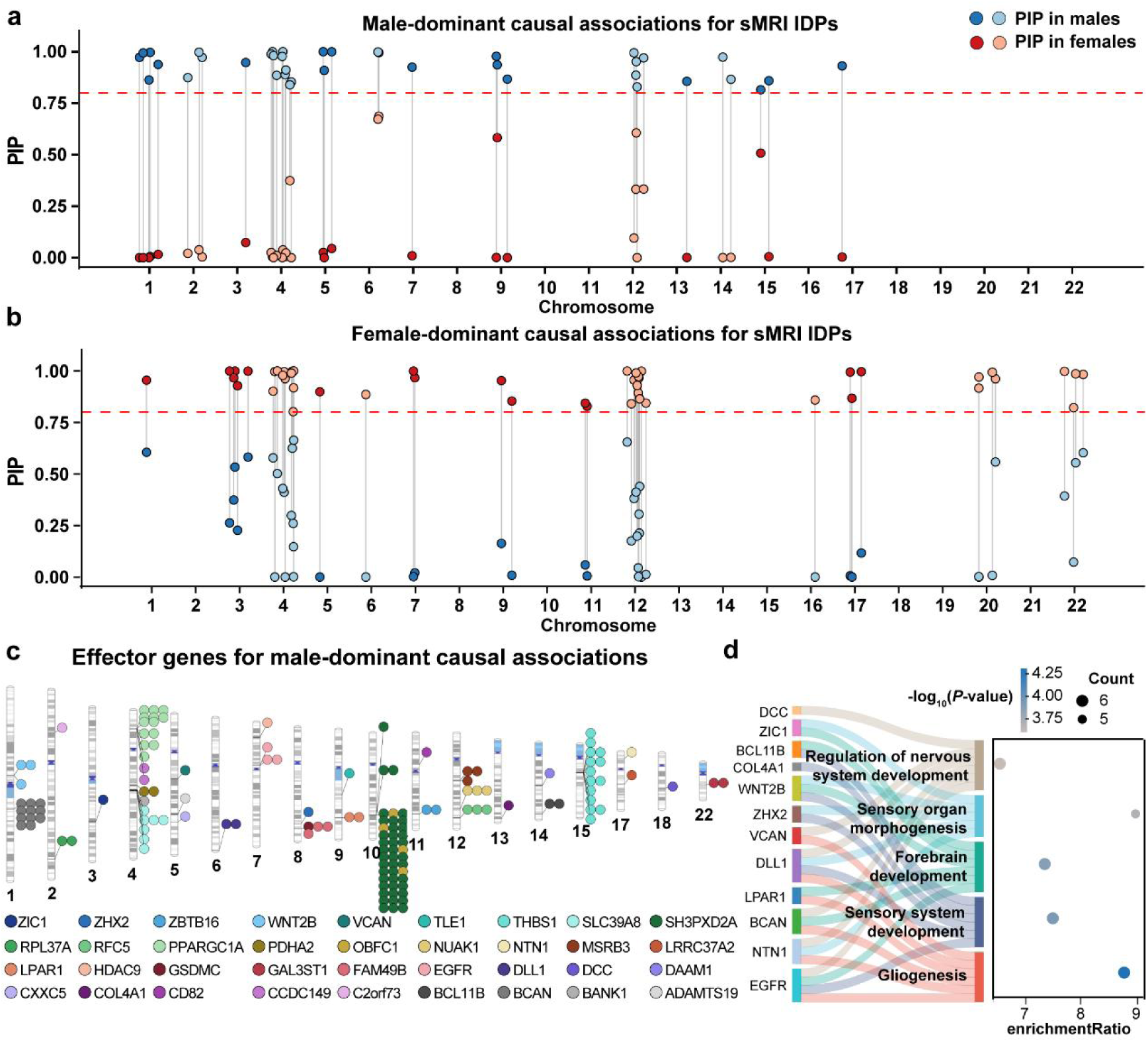
Statistical fine-mapping, gene prioritization, and pathway enrichment. **a, b,** Male-dominant (**a**) and female-dominant (**b**) causal associations for sMRI-IDPs from statistical fine-mapping. Each pair of connected points represents a variant-trait association with a PIP > 0.8 in one sex and an absolute PIP difference (ΔPIP) between sexes exceeding 0.3. Colors distinguish PIP in males (blue) and in females (red). The dashed red line indicates PIP = 0.8. **c,** Ideogram demonstrates the genomic locations of 36 effector genes (colors) identified from 133 male-dominant causal associations (points) using FLAMES with a cumulative precision of >75%. **d**, Enriched biological processes of the 36 effector genes from male-dominant causal associations (q < 0.05, Benjamini-Hochberg FDR corrected). The left column lists the involved genes, the middle column lists the enriched terms, and the right box shows the enrichments. In the right box, the x-axis displays the enrichment fold for each term, point color indicates significance (-log_10_P) of each term, and point size denotes the number of genes overlapping with each term.

**Extended Data Fig. 7.**
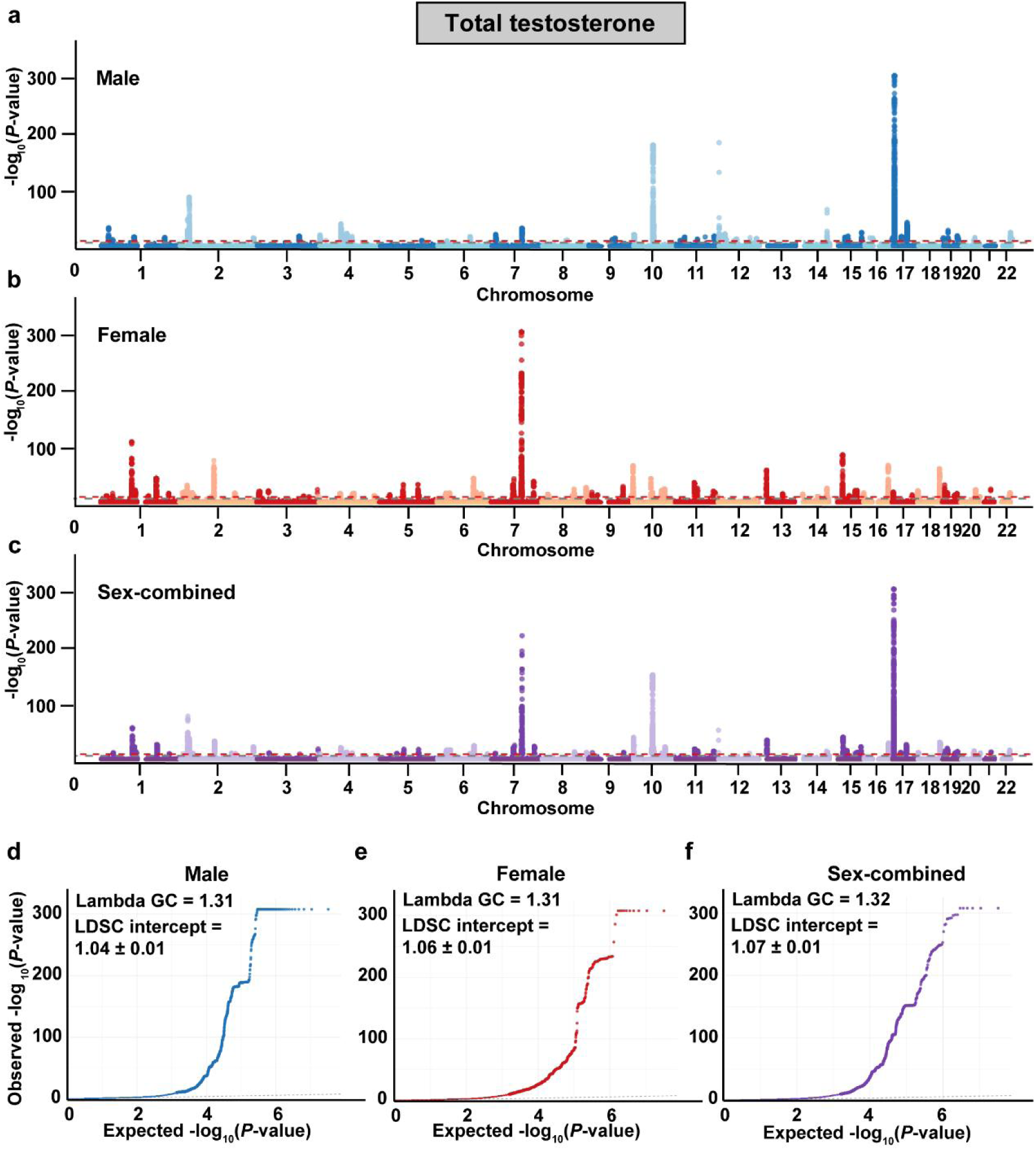
Manhattan and quantile–quantile (Q–Q) plots for GWASs of total testosterone (TT) in the sensitivity analysis. **a-c,** Manhattan plots illustrate genome-wide associations for TT levels in males (**a**), females (**b**), and sex-combined (**c**) samples. The x-axis represents chromosomal position, and the y-axis indicates the strength of association (-log_10_P). The gray and red dashed lines indicate significance thresholds of *P* = 5 × 10^-8^ and *P* = 6.21 × 10^-11^, respectively. **d-f,** Q–Q plots display observed versus expected -log_10_P values for TT-GWASs in males (**d**), females (**e**), and sex-combined (**f**) samples. The genomic control inflation factor (λ_GC_ ) and the LD score regression (LDSC) intercept (mean ± SE) indicate no evidence of population stratification.

**Extended Data Fig. 8.**
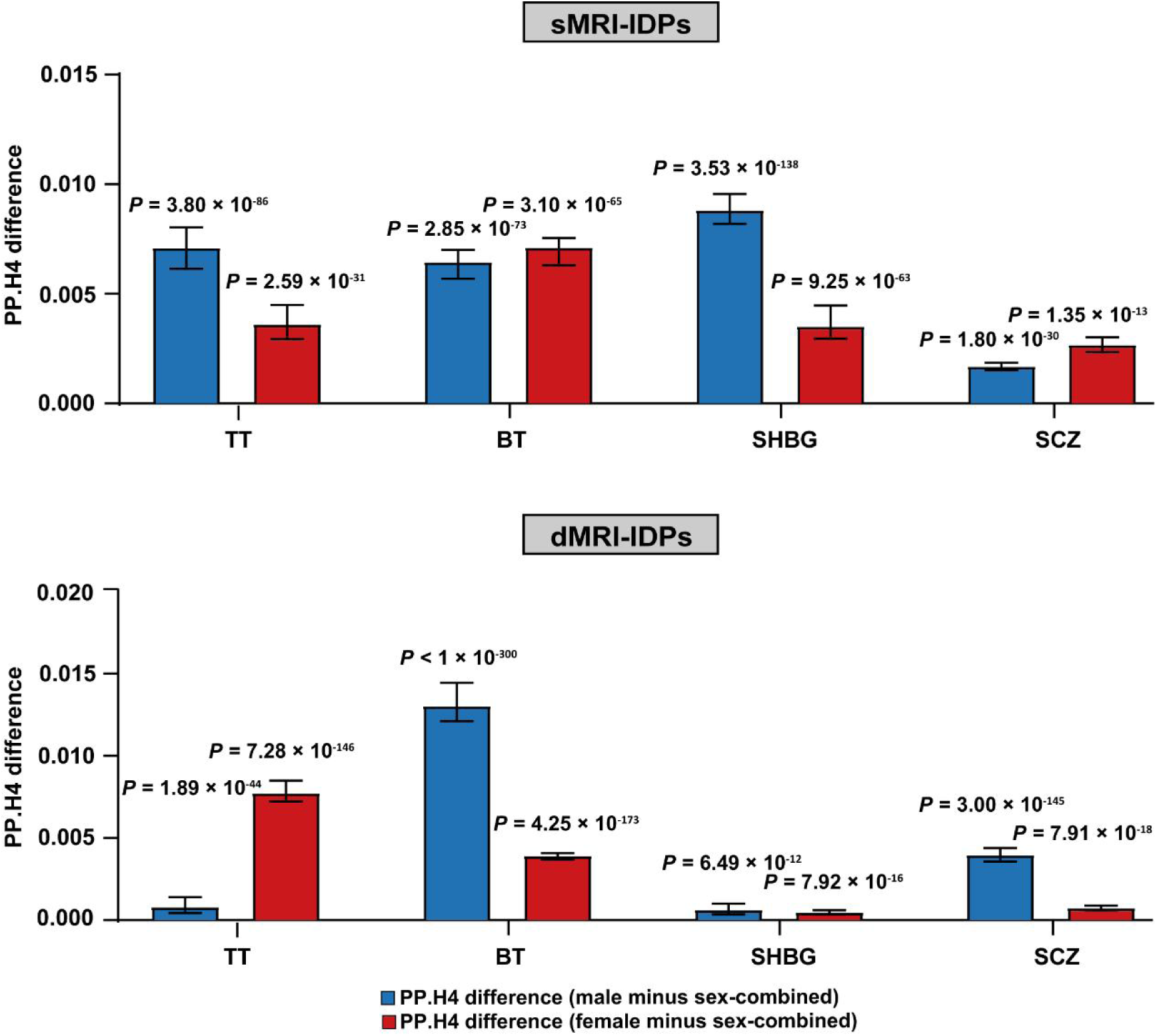
PP.H4 differences of sMRI-IDPs and dMRI-IDPs between sex-stratified and sex-combined analyses. PP.H4 differences (sex-stratified minus sex-combined) for colocalizations of sMRI-IDPs (top) and dMRI-IDPs (bottom) with total testosterone (TT), bioavailable testosterone (BT), sex hormone-binding globulin (SHBG), and schizophrenia (SCZ). Blue indicates male and red indicates female. Values are presented as median with 95% confidence interval (CI), while *P*-values are derived from paired Wilcoxon signed-rank tests. Sex-stratified analyses show higher PP.H4 across all phenotypes compared with sex-combined analyses.

